# An updated evolutionary history and taxonomy of *Mycobacterium tuberculosis* lineage 5, also called *M. africanum*

**DOI:** 10.1101/2022.11.21.517336

**Authors:** Muhammed Rabiu Sahal, Gaetan Senelle, Kevin La, Barbara Molina-Moya, Jose Dominguez, Tukur Panda, Emmanuelle Cambau, Guislaine Refregier, Christophe Sola, Christophe Guyeux

**Affiliations:** FEMTO-ST Institute, UMR 6174 CNRS, DISC Computer Science Department, Univ. Bourgogne Franche-Comté (UBFC), 16 Route de Gray, 25000 Besançon; INSERM-Université Paris-Cité, IAME Laboratory, UMR1137; Université Paris-Saclay, 91190, Saint-Aubin, France; AP-HP, GHU Nord site Bichat, Service de mycobactériologie spécialisée et de référence; Servei de Microbiologia, Hospital Universitari Germans Trias i Pujol, Institut d’Investigació Germans Trias i Pujol, Universitat Autònoma de Barcelona, Badalona, Spain CIBER Enfermedades Respiratorias (CIBERES), Instituto de Salud Carlos III, Madrid, Spain; KNCV Tuberculosis Foundation, Nigeria; Ecologie Systematique Evolution, Université Paris-Saclay, CNRS, AgroParisTech, UMR ESE, 91405, Orsay, France

**Author notes:** **Corresponding author and email address**, Christophe Sola, ^2^Université Paris-Saclay, 91190, Saint-Aubin, France. These two authors contributed equally to the study (Muhammed Rabiu Sahal, Gaetan Senelle). **Repositories**: The search was done in the TB-Annotator 15901 genomes version which is available at: http:// (to be added). The new sequenced genomes are available via NCBI Bioproject accession number: (to be added).

**Keywords:** Mycobacterium tuberculosis complex, L5, Mycobacterium africanum, Evolutionary Genomics, Phylogeography

## Abstract

Contrarily to other lineages such as L2 and L4, there are still scarce whole-genome-sequence data on L5-L6 MTBC clinical isolates in public genomes repositories. Recent results suggest a high complexity of L5 history in Africa. It is of importance for an adequate assessment of TB infection in Africa, that is still related to the presence of L5-L6 MTBC strains. This study reports a significant improvement of our knowledge of L5 diversity, phylogeographical history, and global population structure of *Mycobacterium africanum* L5. To achieve this aim, we sequenced new clinical isolates from Northern Nigeria and from proprietary collections, and used a new powerful bioinformatical pipeline, *TB-Annotator* that explores not only the shared SNPs but also shared missing genes, identical IS*6110* insertion sites and shared regions of deletion. This study using both newly sequenced genomes and available public genomes allows to describe new L5 sublineages. We report that the MTBC L5 tree is made-up of at least 12 sublineages from which 6 are new descriptions. We confront our new classification to the most recent published one and suggest new naming for the discovered sublineages. Finally, we discuss the phylogeographical specificity of sublineages 5.1 and sublineage 5.2 and suggest a new hypothesis of L5-L6 emergence in Africa.

**Impact statement:** Recent studies on *Mycobacterium africanum* (L5-L6-L9 of MTBC) genomic diversity and its evolution in Africa discovered three new lineages of the *Mycobacterium tuberculosis* complex (MTBC) in the last ten years (L7, L8, L9). These discoveries are symptomatic of the delay in characterizing the diversity of the MTBC on the African continent. Another understudied part of MTBC diversity is the intra-lineage diversity of L5 and L6. This study unravels an hidden diversity of L5 in Africa and provides a more exhaustive description of specific genetic features of each sublineage by using a proprietary “*TB-Annotator*” pipeline. Furthermore, we identify different phylogeographical localization trends between L5.1 and L5.2, suggesting different histories. Our results suggest that a better understanding of the spatiotemporal dynamics of MTBC in Africa absolutely requires a large sampling effort and powerful tools to dig into the retrieved diversity.

**Data summary:** *[A section describing all supporting external data, software or code, including the DOI(s) and/or accession numbers(s), and the associated URL. If no data was generated or reused in the research, please state this.]*

The search was done in the TB-Annotator 15901 genomes version which is available at: http://(to be added). The new sequenced genomes are available via NCBI Bioproject accession number: (*to be added*). The authors confirm all supporting data, code and protocols have been provided within the article or through supplementary data files.

## Introduction

*Mycobacterium africanum* is a human and animal pathogenic species that belongs to the *Mycobacterium tuberculosis* complex (MTBC) and that was identified for the first time in 1968 (Castets et al., 1968). It was rapidly shown to present intermediate phenotypic characteristics in between human and bovis tubercle bacilli (David et al., 1978), and later split into two highly related phylogenetic branches, L5 and L6 with specific geographical distribution (Comas et al., 2013). L5 is also known as *M. africanum* West african 1 (MAF1) and as AFRI2-AFRI3 regarding its spoligotype pattern, whereas L6 is known as *M. africanum* West african 2 (MAF2) and as AFRI1 (Comas et al., 2009). L5 is more prevalent in the gulf of Guinea (Ghana, Benin, Nigeria), whereas L6 is more prevalent in the Western part of West Africa (Senegal, Gambia) (de Jong et al., 2009; Gehre et al., 2013). Recently, two important papers pointed the undersampling of L5 and L6 in public MTBC genomes database and laid the foundation for a first sub-classification and diversification history (Ates et al., 2018; Coscolla et al., 2021). It is hypothesized that L5 and L6 could have been endemic to different specific african populations before european colonization, and that these genetic variants could have been less pathogenic than the clinical isolates of the L4 lineage, due to a longer history of co-evolution with humans. Specific L4 sublineages are today highly prevalent in Africa, e.g. L4.6.1 “Uganda”, L4.6.2 “Cameroun”, L4.7 “Congo”, and other modern strains such as L4.8 (Asante-Poku et al., 2015; Malm et al., 2017; Niobe-Eyangoh et al., 2003; Wamala et al., 2015). Hence, the differential history of L4-L5-L6 Africa is of great importance to study the global history of TB. A recent national sampling of MTBC positive Ziehl-Neelsen slides in Nigeria pointed to a very high prevalence of L5 in the south/south-east states of Nigeria (Molina-Moya et al., 2018b). This dense region is geographically close to the cradle of agriculture in Africa, around South-east Nigeria and Cameroon (Molina-Moya et al., 2018b; Phillipson, 1975). It is widely accepted that, once agriculture emerged, the feeding of animals was the first goal of early African growers to maintain foraging and a nomadic way of life. In Africa, and contrarily to the rest of the world, cattle predominated over crops in the food production process. This might have given more time and more chances for animal diseases to spread into human populations, in relation to slow the known environmental changes, such as the ones studied in North-East Africa, and to a slow demographic expansion, slowly accompanied by the Bantu agricultural expansion (Kuper and Kropelin, 2006).

Three main issues are found when dealing with L5 phylogenetics. The first one is undoubtedly the poor sampling and hence the poor statistical representativity of the population studied (Ates et al., 2018). The second one is the important geographical bias that is found in data content in databases, since some demographically important countries and populations, e.g. Nigeria, are only weakly characterized whereas less populated countries, yet more developed, such as Ghana for example, or more recently Ethiopia, start to be well characterized. The third issue is the potential biases and loss of information in evolutionary studies when genome characterization is inferred solely using H37Rv as the reference genome (Sanoussi et al., 2021).

Nevertheless, even with these limitations, some important results were obtained recently. Firstly it is suggested that L5 and L6 emerged in two different ecological niches (Asante-Poku et al., 2016; Comas et al., 2013; Otchere et al., 2018). Whereas L5 is strictly human, L6 is closer to *M. bovis* (Otchere et al., 2018). Another important result is the finding of a new L9 lineage, that is found in East Africa, and the finding of a complex history of *Mycobacterium africanum* (Coscolla et al., 2021). In this last study, Coscolla *et al*. were not too precise in the definition of some sublineages of L5, for example by directly jumping to sub-sublineages of L5.1, thus showing their difficulties to find adequate markers to define hierarchical levels (Coscolla et al., 2021). In another perspective, Sanoussi *et al*. sequenced three new L5 genomes (from Gambia, Benin and Nigeria) using the zero-mode wave guide technology (PacBio®, i.e. producing long read sequencing), and they performed *de novo* assemblies of these new genomes. They showed that classical genome comparison to H37Rv failed to detect some potentially physiologically important genes, a strong limitation in comparative genomics studies that usually tend to limit their analysis to SNPs variation (Sanoussi et al., 2021). More precisely, the pangenome of L5 and L6 is likely to differ from the one of other MTBC lineages. The gene content variation could be larger than expected initially. In addition, gene expression impacting metabolism and pathogenic power is likely to provide different metabolic specificities in L5 versus other lineages (Sanoussi et al., 2021; Yimer et al., 2020).

For all these reasons, we undertook a new study using Whole-genome Sequencing (WGS) of newly recruited L5 and L6 clinical isolates from an african country, Nigeria, as well as from archives from previous studies. We analyzed the genome reads using a new proprietary informatic pipeline, *TB-Annotator*, in order to perform an *in-depth* phylogenetical analysis of this larger collection (Senelle *et al*. manuscript in preparation). We report in this article the description of an improved L5 phylogenetic tree with detailed evolutionary events list for each branch, and we suggest new hypotheses regarding its history.

## Methods

### Origin of clinical isolates of newly sequenced L5-L6 genomes

The 25 newly sequenced *M.africanum* (numbered CUS0000001-CUS0000025) were recovered from well-characterized human TB cases from archive DNAfrom previous published studies whose DNA had previously been sent between 2007 to 2019 to the Infection Genetics and Evolution of Emerging Pathogens research team (IGEPE, Institute of Integrative Cell Biology, Gif sur Yvette) (Lawson et al., 2012; Molina-Moya et al., 2018a; Molina-Moya et al., 2018b). Another origin of samples is culture positive L5-L6 isolates from the French national TB reference-associated laboratory in Bichat Hospital Paris, France. Altogether, these included isolates recovered from Nigeria (n=13), and France (n=12) with 11 patients representing seven West-African countries nationalities and one isolate representing a patient of French nationality (data available upon request).

The selected samples were initially screened as L5-L6 based on a simple MIRU24 locus copy number assay, and further confirmation by a Line-probe PCR-based assay (Hain Lifescience GmbH-Brücker Molecular diagnostics, Gehren, Germany). CUS0000001-09 and CUS0000022-25 are archive DNA from Nigeria whereas CUS0000010-21 are L5-L6 isolates from Bichat Hopital. CUS0000001 and 0000008 represent isolates from Lawson *et al*. 2012 collection, while the remaining eleven are from collections either recovered from Molina-Moya *et al*., 2018a or 2018b collections. In this study, newly recruited L5-L6 DNAs from Nigeria were selected from patients with a microbiological diagnosis of TB characterized by the presence of 2 copies of MIRU24 as assessed by MIRU-VNTR analysis. The demographic data (age, sex, country of birth, years of diagnosis especially applicable to France isolates) and clinical (location of disease, sputum smear status, and previous diagnosis of TB) characteristics of the patients, when available will be reported elsewhere.

### Culture, DNA purification methods, and Whole-Genome Sequencing on Illumina

MTBC clinical isolates were firstly inactivated in P3 laboratory from Coletsos solid media grown cultures. DNA was extracted using DNeasy® UltraClean 96 Microbial Kit (Qiagen®, Hilden, Germany). A Nanodrop® device (ND-ONE-W) was used to control purity and concentration for each sample (Thermofisher®, Waltham, USA). The Qiagen® kit allows to obtain good quality purified DNA as assessed by a 280/260 and 260/230 ratio close to 2 and the concentration of the DNA was sufficient to perform library preparation (concentration ~100 ng/μl). Library was produced using the Nextera® XT DNA Library Preparation kit (Illumina, San Diego, USA)-with a control of the fragment size on a 2100 Bioanalyzer® Instrument (Thermofisher®, Waltham, USA). Sequencing was performed using the Miniseq® High Output Kit (Illumina®, San Diego, USA) generating paired-end reads of 150 bp. Quality of reads was then analyzed by *fastqc* (Andrews, 2010).

### Variant definition and presentation of the *TB-Annotator* and *CRISPR-builder-TB* pipelines

Figure 1 shows the general algorithm that was used to process the Short Read Archives (SRA) produced in house or recovered in public repositories. A brief description of the pipeline can be found in (Guyeux et al., 2022). At the time of writing, the V1 database contained 15909 SRA; results were also checked in part using the expanded V2 version (Senelle *et al*. in preparation).

**Figure 1:**
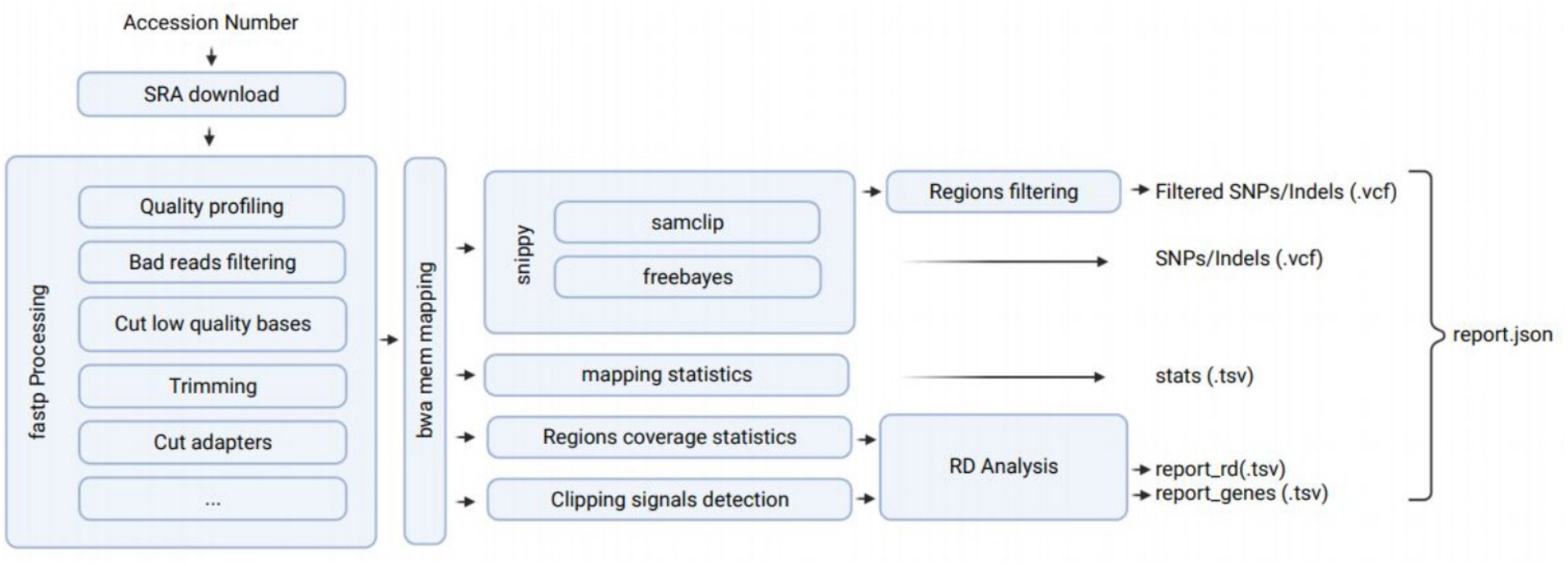
General algorithm

### Selection of samples from the Public databases for uploading on TB-annotator platform

NCBI SRA Projects from all studies known or likely to include L5 or L6 samples were incorporated in TB annotator platform using the keywords “*M. africanum* ; *Mycobacterium africanum* ; *Mycobacterium tuberculosis* L5 ; *Mycobacterium tuberculosis* L6”.

### Sample selection for the study, TB-annotator platform, and tree reconstruction

All samples sharing the defining SNP shared by L5 samples were selected; as L5 defining SNP we chose the C9566T SNP (selection of 360 isolates with 133 exclusive variants and 548 more at >95% specificity) instead of C1799921A (selection of 359 isolates with only one exclusive variants and 676 variants at >95% specificity. They confirmed to be monophyletic in the visualization tool. Several subselections were performed to further explore L5 intra-lineage diversity. Pairwise distances between clusters were systematically computed for the defined clades in order to asses the epidemic (< 15 *SNP*) versus historic (>15 SNP) status. Because of complexity issues, this is done only for clusters of size (< 200 isolates). The unrooted tree was built using a similar approach as the one published in (Guyeux et al., 2022). Briefly, we used RAxML on a multi fasta-file produced using *TB-Annotator*; our files initially contained 15901 SRAs at start, the bincat model was used instead of the GTR model and no outgroup was chosen (S t a m a t a k i s, 2 0 1 4). A confirmation of the results using the extended *TB-Annotator* v2.1 version 85000 SRA, that unfortunately was not much richer in L5, was obtained (results not shown). Variants were searched in *TB-Annotator* v1 or v2.1 based on SNP position or using the SPDI Model, with H37Rv as the reference genome (NC_000962.3 reference sequence) (Holmes et al., 2020).

## Results

### 3.1 Identification of new L5 sublineages

A first unrooted tree of L5, shown in **Figure 2**, was obtained on an alignment analyzing 360 L5 samples. This first tree was obtained using not only SNPs but also presence of IS*6110* at the same position, shared deletions and shared absence of H37Rv genes. This original combined sample requires to use binary info of shared(1)/unshared(0) patterns.

**Figure 2:**
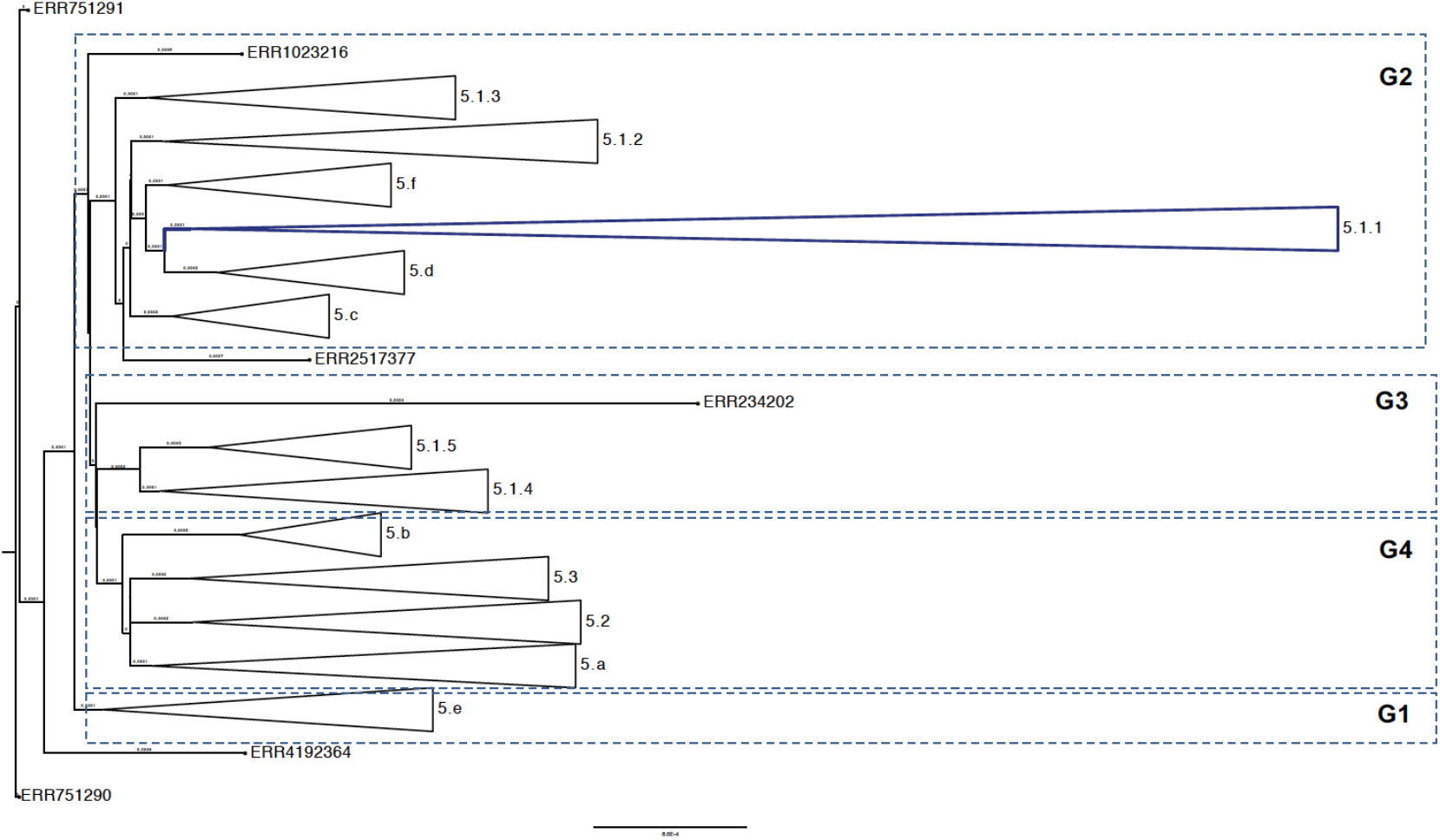
Unrooted tree of L5 samples defining main phylogenetic groups

We first confirm the overall predominance of the L5.1 sublineage in the current sampling as described by Coscolla et al (2021). Second, we confirm the existence of other L5 sublineages such as L5.2 and L5.3 (Coscolla et al., 2021).We identified 6 new subclades, first identified as 5.a to 5.f. Some of them grouped with L5.1 main sublineage, namely those transiently named 5.c, 5.d and 5.f. The subgroup including them and former 5.1.1, 5.1.2 and 5.1.3, supported by 30 SNPs, is denoted G2 below. The third main monophyletic group named G3 consists of subclades 5.1.4 and 5.1.5, supported by 126 SNP, and the last main monophyletic group, G4, supported by 72 SNPs, includes 5.2, 5.3 and two new subclades 5.a and 5.b.The new clade transiently denoted 5.e, at the bottom of the tree on **Figure 2** (also noted G1 thereafter), seems basal but this is due to the fact that the tree was unrooted (see below for a description of its relative position in L5 diversity).

On the one hand, 5.e seems to have a basal position. On the other hand, when we look at the arrangement of the 3 other groups, it appears as shown in Figure 2 that 5.1.4. and 5.1.5 do not clade with 5.1.1, 5.1.2 and 5.1.3, but rather with 5.2 and 5.3, which would argue against the current nomenclature and scheme proposed in (Coscolla et al., 2021). However, the position of the ancestral nodes of these three groups is rather confusing, and this is reinforced by the weakness of the supports of the branches near the root (data not shown). The phylogeny does not seem to be resolved in view of the very short length of the branches there: as often in such cases, the further back in the tree one goes, the more the phylogenetic relationship becomes blurred. Before going into more detail on the new clades and sublineages definition, we will first investigate this lack of resolution at the base of the tree, and then propose a new phylogeny of L5.

### 3.2 Genetic features characterizing the main L5 large monophyletic groups

We analyzed the G1 to G4 groups by pair and by triplet, to see which tuples had similar mutations, a sign of a common ancestor. G1 and G2 have 4 exclusive SNP variants at 97.64%, meaning that all the strains of this pair have these 4 mutations (C558750T, G4313357A, T1407273A, and G4343653A), while we can find 5 strains outside with these mutations. Conversely, the other pairs of strains shared nothing in common. At the triplet level, (G1, G2, G3) shared 10 exclusive SNP variants, and 19 more SNP variants with exclusivity being larger than 95%, while no other triplet share any other exclusive element.

We took into account that only (G1, G2) on the one hand, and (G1, G2, G3) on the other hand, emerge from our exclusivity study to build the tree detailed in **Figure 3** based on **Figure 2**. **Figure 4** presents the same tree, however following a hierarchical naming process.

**Figure 3:**
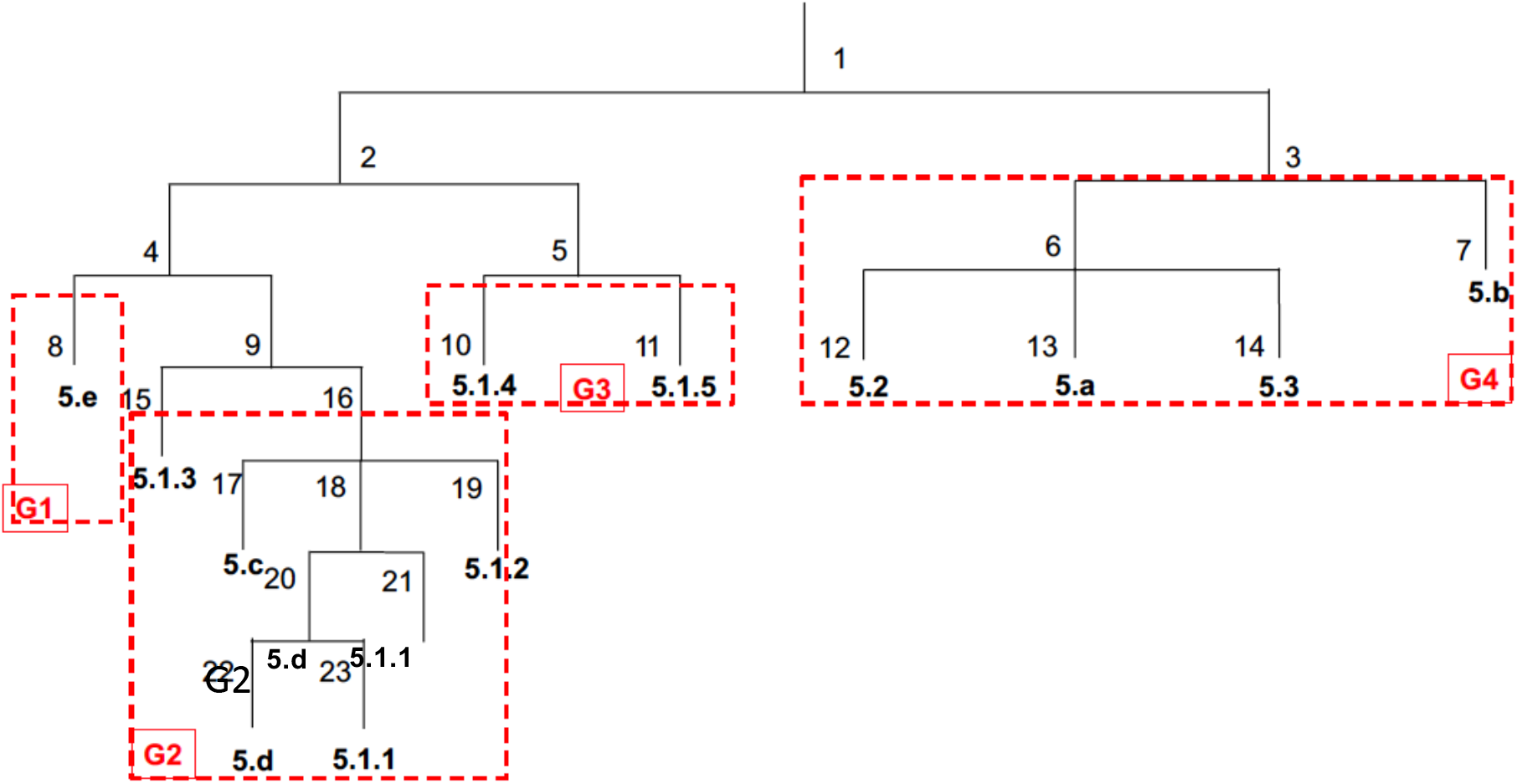
Consensus phylogenetic relationships between reference sublineage naming (Coscolla et al., 2021) and transient naming of new clades (this study) (new 5.3 not present at this stage)

**Figure 4:**
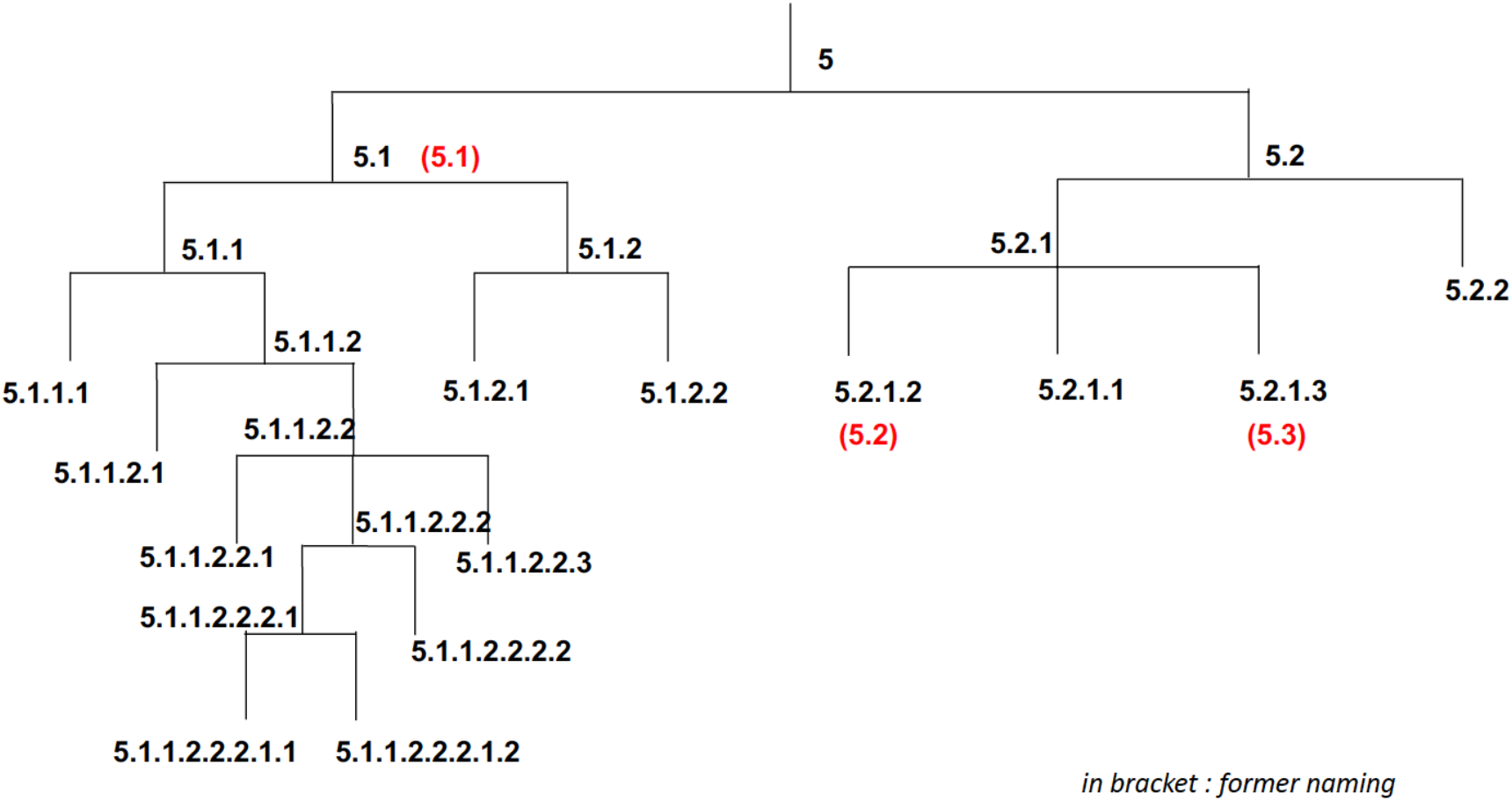
Consensus phylogenetic relationships with new hierarchical level naming and reminders of main correspondence with M. Coscolla *et al*. classification (under brackets) (new 5.3 is not present at this stage)

In this phylogeny proposal, the only internal node not having its position proven through the data is the node giving rise to L5.2, whose resolution may still be problematic. However, we will see in the CRISPR locus structural study section of this article that a recombination of IS*6110* upstream of spacer 35 is a specific feature of 5.1, which is never present in 5.2, and this fact strengthens our proposal. Moreover, this tree is in good accordance with that of Coscolla *et al*. (2021). In the following, we will present the elements supporting the different nodes of the proposed tree.

### 3.3 Genetic features characteristic of the main sublineages

Our first objective is to provide support for our proposed L5 tree, as depicted in **Figure 4**. We describe below the specificity of each node (internal or leaf) starting from its root, on a sample of 359 and not 360 since ERR4872200 does not contain any of the 5.1, 5.2, and 5.3 defining SNPs. The classification of this SRA remains pending. The L5 lineage has 133 exclusive SNP variants of its own, and 548 variants with more than 95 % exclusivity, (cf. **Supplementary Material 2**). A SNP in *gyrA* for example, very well defines L5, namely C9566T (n=360). The absence of three specific L5 genes, Rv2478c, Rv1978, and Rv1979c also defines L5. Two isolates, ERR439931 and ERR4872094 over 360 did not lack Rv2478c. Alternatively the lack of Rv2478c is found in >360 isolates in the *TB-Annotator* database, among which ancestral L2 from Kwa-Zulu Natal (SRR10315503). Concerning the known region of differences, RD713 (Rv1977-Rv1979c) is common to all L5 (Mostowy et al., 2004). A search of lack of Rv1978 in *TB-Annotator* retrieves 390 SRAs, among which a few L2, L3, L6 and animal isolates (Mostowy et al., 2004). Last but not least, 350 strains (97.49% of L5) have an IS*6110* at position 4185790, while only 10 out of 15542 not-L5 isolates distributed throughout the tree have this IS too. To put it in a nutshell, 133 SNPs define this lineage (with 548 exclusive >95% variants), these other characters being listed as almost exclusive. Let us go further into the tree.

#### First sublevel

Internal node 5.1 is defined by T3452C, and is constituted by 278 isolates. In addition to its 10 exclusive SNP variants (plus 19 at > 95% exclusivity), we can note the systematic absence of 3 genes (Rv1333, Rv1334, and Rv1335), the first two being almost exclusive, corresponding to RD711 as region of difference (10 isolates too outside this selection) (Mostowy et al., 2004). Section 3.5 on CRISPR will provide another characteristic of this lineage. Of note, in this L5.1 as well as in L5.2 and L5.3 nodes, we identify more reliable subgroups than in the previous nomenclature. This requires a renaming of all groups, which is detailed below and with a correspondence with former publication as shown in **Table 1**.

**Table 1:**
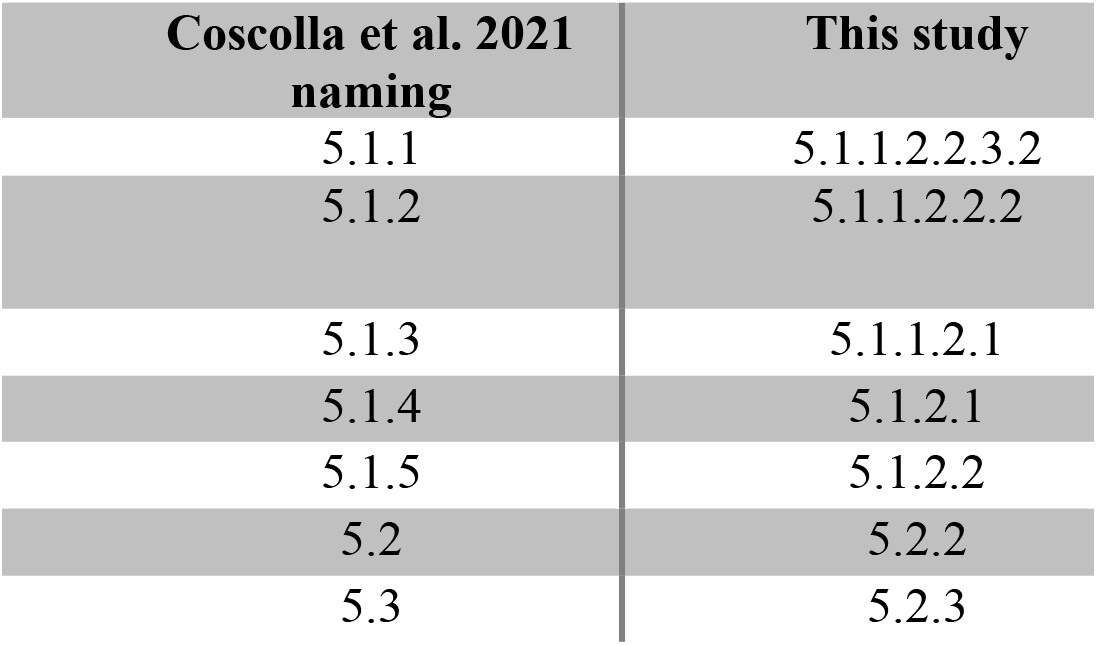
conversion table between the reference nomenclature and our new naming

| Coscolla et al. 2021<br>naming | This study |
| --- | --- |
| 5.1.1 | 5.1.1.2.2.3.2 |
| 5.1.2 | 5.1.1.2.2.2 |
| 5.1.3 | 5.1.1.2.1 |
| 5.1.4 | 5.1.2.1 |
| 5.1.5 | 5.1.2.2 |
| 5.2 | 5.2.2 |
| 5.3 | 5.2.3 |
| **5.1.1** | **5.1.1.2.2.3.2** |
| **5.1.2** | **5.1.1.2.2.2** |
| **5.1.3** | **5.1.1.2.1** |
| **5.1.4** | **5.1.2.1** |
| **5.1.5** | **5.1.2.2** |
| **5.2** | **5.2.2** |
| **5.3** | **5.2.3** |

The node 5.2 encompasses the lineages 5.2.2, 5.2.3 (Coscolla et al., 2021), and two new discovered ones, namely: 5.2.1 and 5.2.4 (**Table 1 and Figure 4**). The 75 strains of L5.2 are defined by exactly one exclusive variant, namely C3471147T (0 at 95%). They have an average of 578 SNPs of difference (standard deviation of 150).

Our last analyses suggest that the L5 tree should include another branch that we propose to name L5.3 (that differs from the former L5.3 from Coscolla *et al*). This new leaf 5.3 completes this first level of the phylogenetic tree. It is constituted by only 6 isolates at present, but could represent an undersampled subbranch of the tree (cf. **Figure 5 and Figure 10**). The 6 known samples share 104 exclusive variants (0 at 95%), one of them being C96339T. Their mean distance is equal to 219, with a standard deviation of 95. This clade was not described in the literature nor in *TB-Profiler* (Napier et al., 2020). Three of these genomes had been previously studied in the largest study performed until now on L5-L6 genomes in Ghana and published in (Otchere et al., 2018).

**Figure 5:**
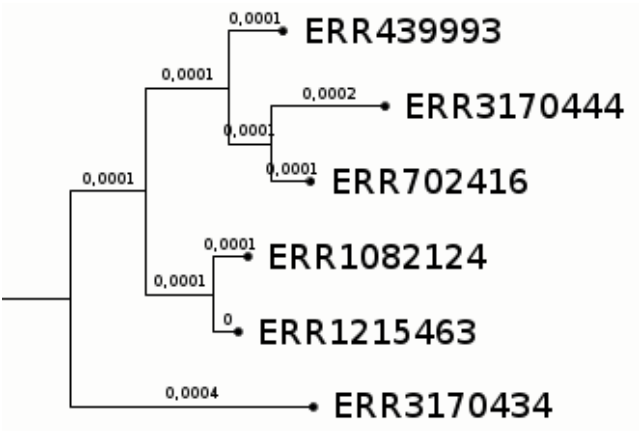
5.3 sublineage

#### Second sublevel, with 5.1 as ancestor

The node 5.1.1 (212 isolates) is the ancestor of lineages 5.1.1.2.2.3.2 (5.1.1), 5.1.1.2.2.4 (5.1.2), and 5.1.1.2.1 (5.1.3), and of a collection of new sublineages (all number in brackets referring to the classification by (Coscolla et al., 2021). This clade, can be defined by the SNP C558750T. It has 2 exclusive SNP variants, and 3 other at 95%. Rv1523 is absent in all these strains, but this is not an exclusive character, as 38 clinical isolates outside the clade share the same deletion.

The node 5.1.2 (65 isolates) is the ancestor of 5.1.2.1 (5.1.4) and 5.1.2.2 (5.1.5) (again all number in brackets referring to (Coscolla et al., 2021)). L5.1.2 is defined by the SNP C118646T and 38 SNP variants are exclusive (and 6 others at 95%). It is constituted by 65 isolates, which are at an average distance of 417 SNPs (standard deviation of 113).

#### Second sublevel, with 5.2 as ancestor

L5.2.1, at the top of the tree in **Figure 6**, is constituted by two isolates only: ERR439960 and ERR439985, both from Benin. These two isolates are far from each other (108 SNPs of difference), and far from all other strains, with 152 exclusive variants (0 > 95%), one of them being C15558A. Furthermore, they share an IS*6110* at position 3841699, and a region of difference at position 3738497-3738697 (200 bp).

**Figure 6:**
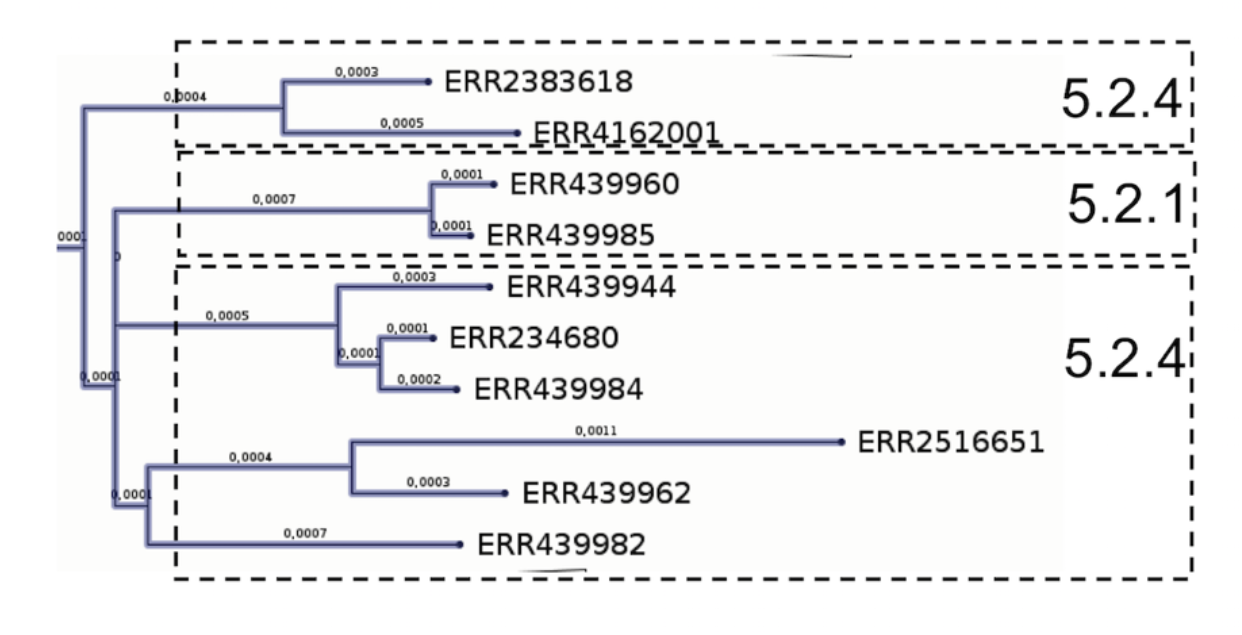
5.2.1 and 5.2.4 sublineages

L5.2.2 (5.2 in in (Coscolla et al., 2021)) is constituted by the 36 isolates of **Figure 7** that are defined by G30475A (63 exclusive variants, 0 at 95%). 35 isolates have an exclusive IS*6110* at position 2129642, which is missing in ERR2704812 originating from Cameroon. The mean distance is of 438 SNPs, and the standard deviation is equal to 162.

**Figure 7:**
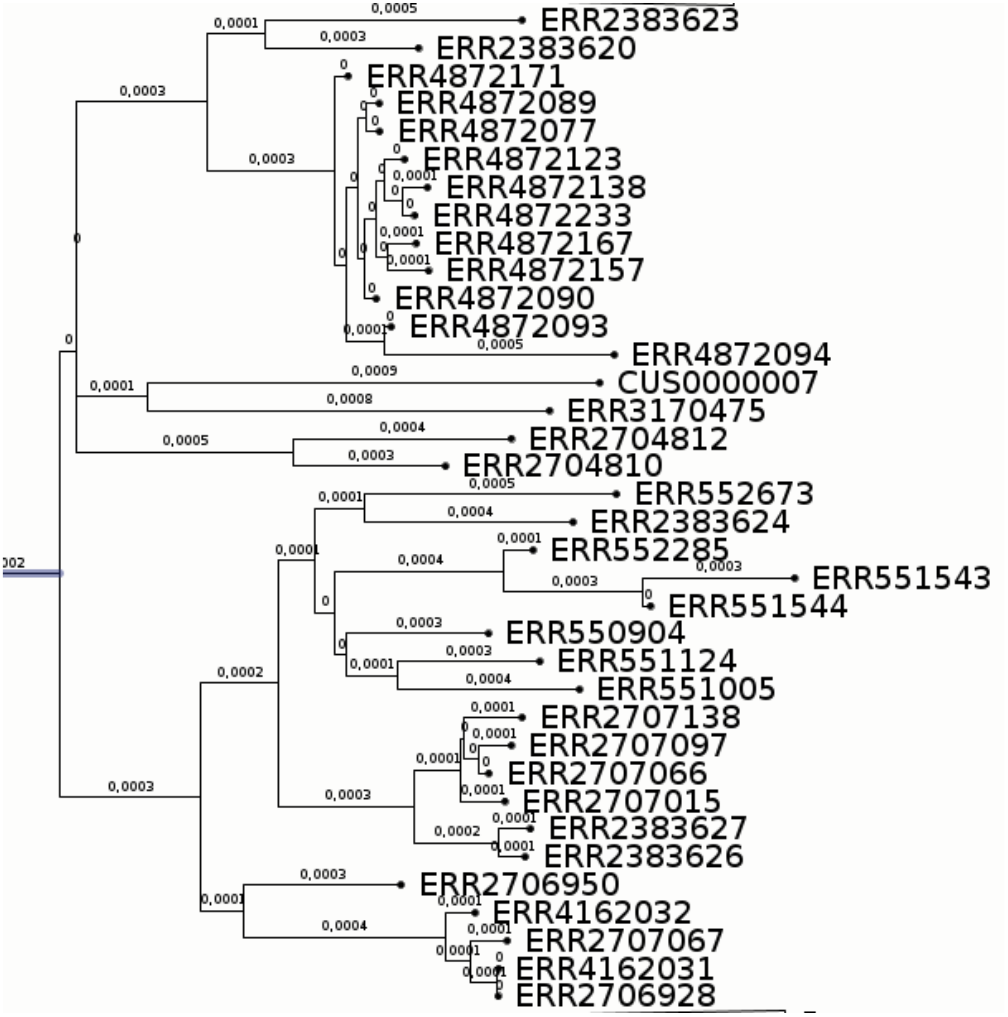
L5.2.2 sublineage or L5.2 in (Coscolla *et al*., 2021)

L5.2.3 (5.3 in (Coscolla et al., 2021)) has 29 isolates defined by A47099T, as shown in **Figure 8**. They share 59 exclusive variants (8 > 95%) and present an average distance of 432 SNPs (standard deviation: 117). L5.2.4, the remainder strains of **Figure 6**, is made-up of 8 isolates. They have two exclusive variants (e.g., G587327A) and a mean difference of 514 SNPs (standard deviation: 124).

**Figure 8:**
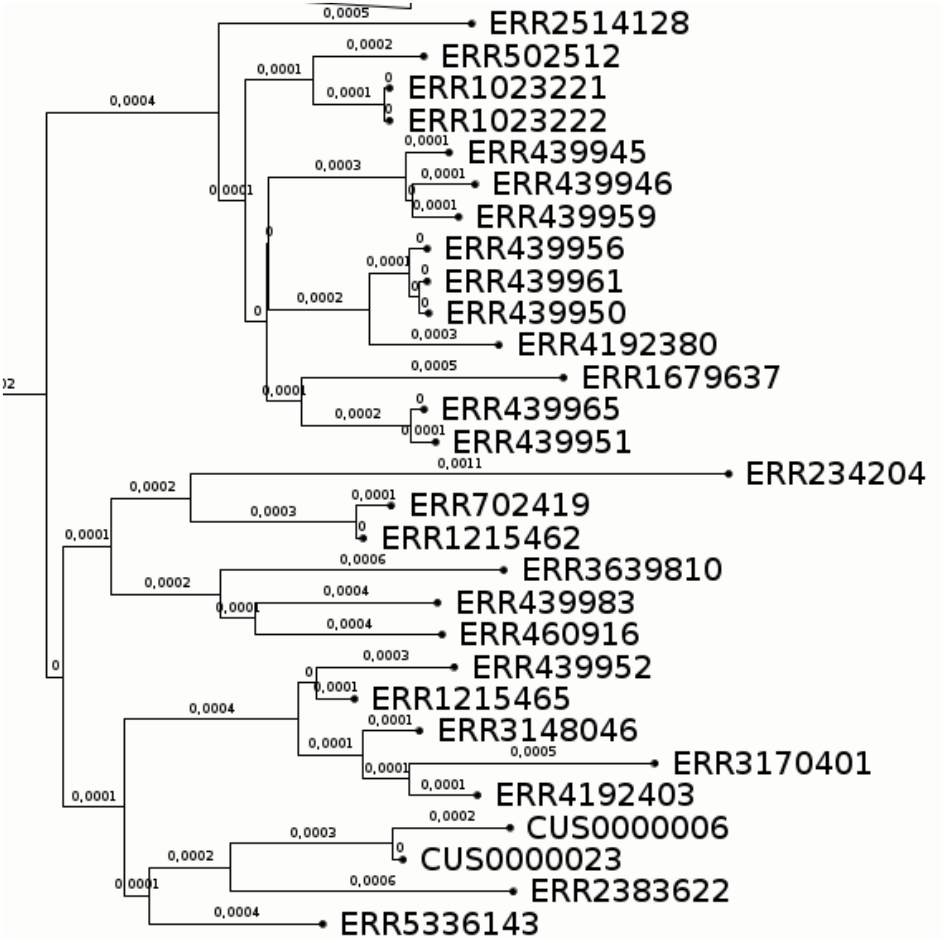
5.2.3 sublineage

### 3.4 A thorough classification of the L5 sub-lineages

Further details of all phylogenetic trees built for L5.1 and L5.2 sublineages are described in detail in the **Supplementary Material 2**. The findings on new phylogenetically relevant SNPs, on SNP distances inside these phylogenetical branches, on SNPs specificity and on CRISPR locus architecture can else be found in the **Supplementary Material 1-4**.

### 3.5 CRISPR locus structure investigation

The CRISPR locus has long be known as a locus evolving by deletion and with good global concordance with SNP-based phylogenies. We describe below its structure in the different L5 sublineages. In what follows, spacer numbers are provided in the extended nomenclature at 68 spacers, whereas SIT there are given to provide the spoligotype value with former 43 spacers format (Couvin et al., 2019; Kamerbeek et al., 1997; van Embden et al., 2000). Lineage 5 is defined by the absence of spacers 18-22 on the one hand, and 51-60 on the other in the 68 spacer format (Brudey et al., 2004); (Refrégier et al., 2020). They all have IS*6110* located upstream of spacer 35. They do not have specific spacer mutants. Unlike the strains of L2, L3 and L4, they do not have DR6 between spacers 65 and 66, but have DR4 and DR5 between spacers 66-67, and 67-68 respectively (Refrégier et al., 2020). In terms of duplications, there is only that of spacer 35 between spacers 41 and 42 to note. And as we will see below, recombinations between IS*6110* are partly responsible for the diversity of spoligotypes in the lineage. As stated previously, all L5.1 isolates have lost their 4 spacers in front of spacer 35 (Ates et al., 2018). It sometimes occurs in L5.1 that the deletion is a bit larger than these 4 spacers. Conversely, these 4 spacers or at least one of them are almost always present in L5.2 and L5.3 (Ates et al., 2018). A further investigation of this specific CRISPR area shows that, during the L5.1 diversification, a recombination between the usual IS*6110* in front of spacer 35, on the one hand, and an IS*6110* after spacer 30, on another hand, has led to the appearance of such a hole at the origin of Lineage 5.1 (Guyeux et al., 2021; Refrégier et al., 2020). As it is hypothesized that the CRISPR locus was acquired once only, by lateral gene transfer from an environmental host, acquired spacers are found only in lineages having predated MTBC last common ancestor, and evolve since by deletion only^1^, and it is not surprising that this hole is found in all descendant lineages of L5.1. Other points that can be made about CRISPR in Lineage 5 include:

- low copies number of IS*6110* in CRISPR, and thus reduced possibilities for evolution in this locus, as in the rest of the genome, this considerably slower the pace of evolution of the CRISPR locus.
- few modifications in the regions bordering the CRISPR (the CRISPR-associated genes *cas* on the left, and the part flanking the locus on the right).

All the characteristics of the CRISPR locus that have produced using “*CRISPR-builder-TB*”, or produced automatically, or collected from other sources (London School of Tropical Medicine, Jody Phelan, personal communication) been gathered for all L5 studied isolates can be found in **Supplementary Material 3**.

### 3.6 SNP discovery and Specificity

When comparing the SNPs informativity level found in this study and the one described in the supplementary material of Coscolla et al. 2021 (n=312), we found an excellent agreement. For some sublineages such as the former L5.1.1 (L5.1.1.2.2.3.2 in this study), the agreement was low (42%), whereas for other subbranches such as former L5.1.3 (L5.1.1.2.1 in this study) the agreement is very high (94% in this case). This is likely to be due to the fact that, given the undersampling of our global dataset, true SNPs discovery is less supported than false SNP discovery for sublineages that are presenting a higher, yet only very poorly investigated, genetic diversity (cf. **Supplementary Material 4**). Since the search for SNPs in *TB-Annotator* was done using a hierarchical process (SNP position followed sometimes by SPDI search), we sometimes unraveled interesting genetic variants. One such example is the case of the finding of three variant alleles in various lineages at position 2187276 (L5.2.3 A:G, L4.1.2 A:T and another L4.1 A:C). Each SNP specificity was verified by individual search in the *TB-Annotator* v1 database and pipeline (see **Supplementary Material 4**). Three other likely convergent but interesting variants detected by Coscolla *et al*. our by this study are SPDI:NC_000962.3:1738750:C:A in *ileS* that is found in both L5.2.2 and in two L3 sublineages, SPDI:NC_000962.3:2361310:C:T in *helZ* that is found in L5.1.12.2.2 and in all L4.5 genomes, and SPDI:NC_000962.3:2433085:G:A in *lppM* that is found in L5.2.2 and in the L2 sublineage L2.2 M5.

Our pipeline is analyzing all genes, without discarding PE-PPE genes nor other repetitive sequences such as Insertions Sequences, however only 82 mutations were found in PE-PPE genes (**Supplementary Material 4**).

To confirm the phylogenetic relationships obtained from binary analyses, we extracted the SNP from 51 samples representative of the L5 diversity (random selection). The corresponding alignement totalled 8701 nucleotides including those belonging to PE-PPE. We built a tree implementing a standard molecular evolution model. We retrieved the same topology as with binary data except for some subbranches of L5.2.4, some of which clustering with L5.2.2 (**Supplementary Material 5**). To confirm that this slight difference did not originate from including PE-PPE genes, we built a second alignment excluding PE-PPE SNPs (**Supplementary Material 5**). The 7840 nucleotides alignment resulted in the same phylogeny except for L5.2.4 subbranches, with a mean 5% increase in branch lengths. This confirms that the proposed classification is highly reliable, with the exception of L5.2.4. This also indicates that PE-PPE SNPs contribute to around 10% of the retrieved diversity and that this diversity adds only little noise to the rest of the phylogenetic information.

### 3.7 Geographical specificity of L5.1 versus L5.2

We tried to search for an eventual geographical specificity of L5.1 and L5.2 by trying to link the taxonomic position of the main sublineages described to their known geographical region of origin (**Figure 9** and **Supplementary Material 3**). We adopted the same country classification as the one found in (Coscolla et al., 2021). West of West Africa includes Gambia, Senegal, Mauritania, Sierra Leone, Liberia, Guinea, Ivory Coast and Mali; East of West Africa includes Ghana, Benin, Niger, Nigeria and Burkina Faso; Central Africa includes Cameroon, Gabon, Congo, Democratic republic of Congo, and Equatorial Guinea. By performing a Fisher exact statistical test to look for the homogeneity of distribution of sublineage 5.1 and L5.2, we suggest a statistical trend that L5.1 and L5.2 are heterogeneously distributed in the three african regions (p-value = 0.07439, close to 0,05). To support this, we provide a map of the distribution of L5.1 and L5.2 in **Figure 9**. We suggest that L5.1 and L5.2 sublineages have different phylogeographic origin and different epidemic histories, as what had already been demonstrated for L6 but not for L5 in (Coscolla et al., 2021). However this result needs to be confirmed in a new study on an more representative and extended dataset.

**Figure 9:**
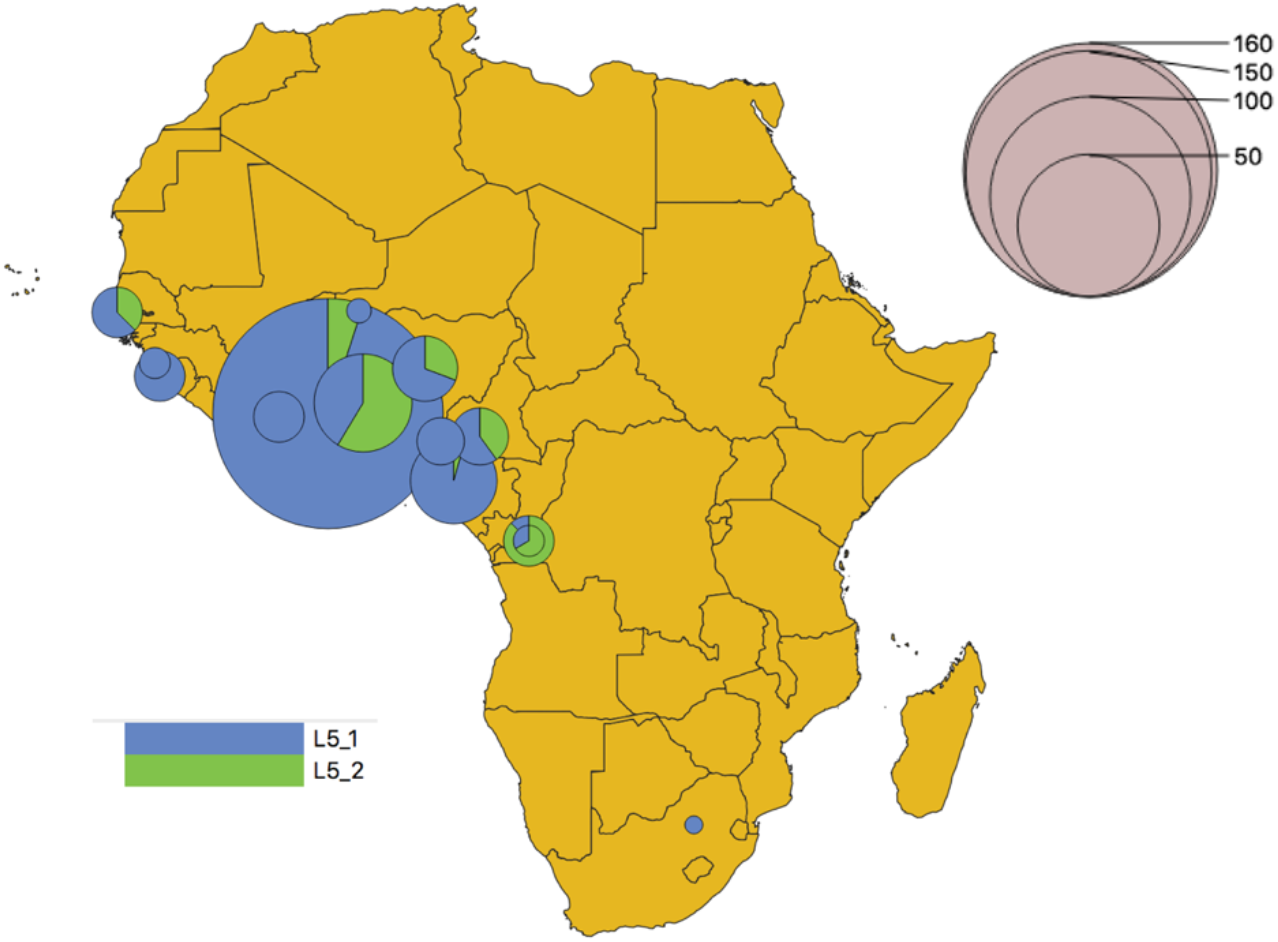
L5.1 and L5.2 (newly defined) geographical distribution

**Figure 10:**
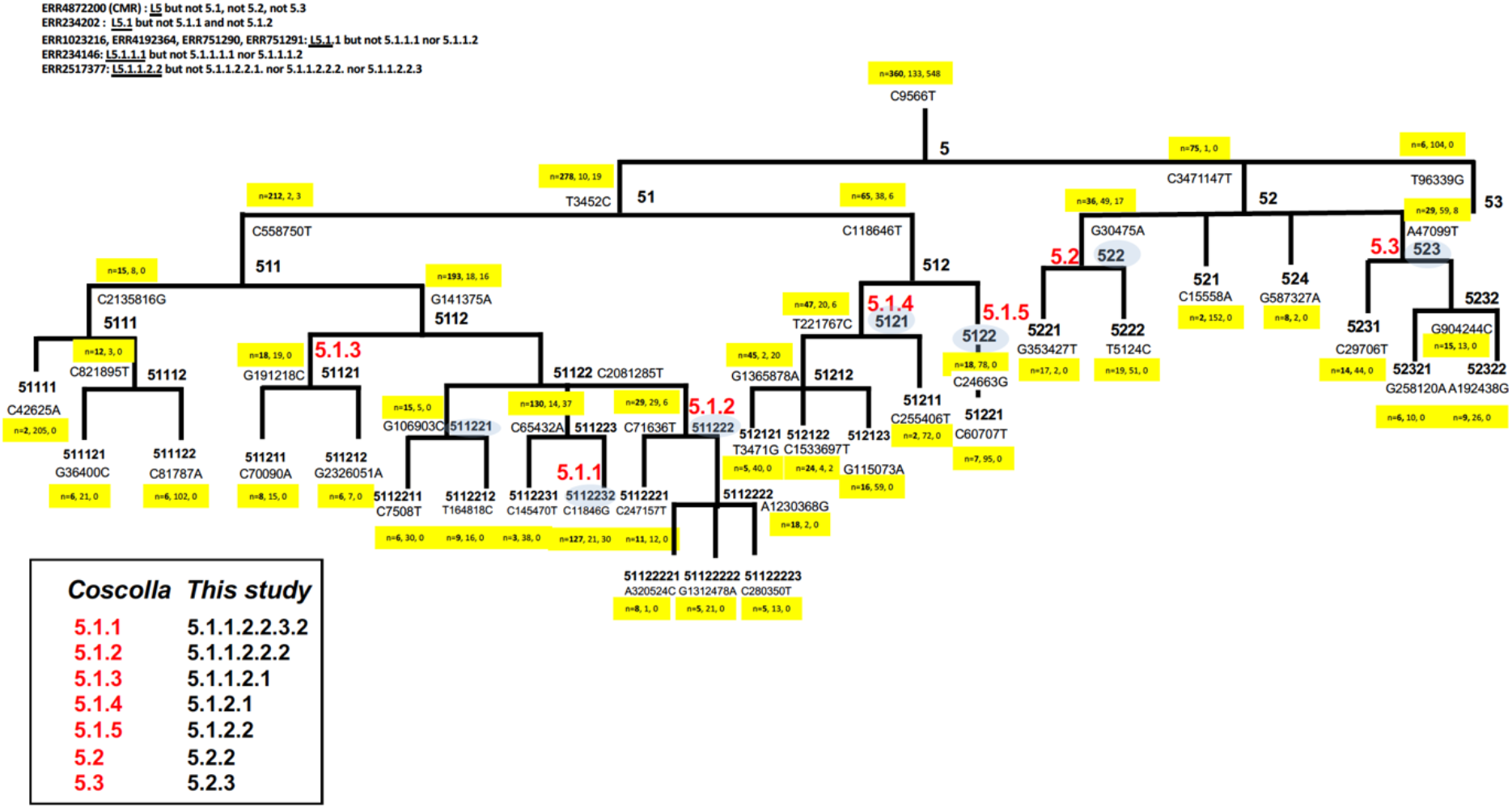
A final phylogeny of Lineage 5 with SNP barcode proposal. *in red : ancient naming (Coscolla et al. 2021); yellow frame: number of genomes in this group, number of defining SNPs (100% exclusive), number of SNPs (>95% exclusive)*; *top left: peculiar isolates IDs that do not fit within this classification*

## 4. Discussion

The reconstruction of the evolutionary history and taxonomy of L5 is important for the understanding of the natural history of tuberculosis in Africa and to predicts its evolution. It is a yet fragmented endeavor since there remains a strong under-representativity of these genomes in public databases. However, as shown in **Figure 10**, that summarizes our results, this study provides a considerable increase in the phylogenetical analysis of L5: (1) a better description of basal groups, (2) the identification of new reliable SNPs at each subbranch (3) and an integrated phylogenetic reconstruction of the entire L5 lineage.

Our findings comfort the recent studies found on L5, namely the predominance, in the present sampling of what we still call L5.1 (Ates et al., 2018; Coscolla et al., 2021).

The main aim of this study was to build an exhaustive collection of representative L5 genomes and to analyze their structure using a combination of bioinformatical analytical pipelines, so called “*TB-Annotator*”. First, let us note that the phylogenetic resolution is more difficult towards the root, which is expected. However it can be solved with the so-called exclusive variants approach. For example, we were able to cleanly define lineage 5.1 (new and old, no modification), which was previously undefined (Coscolla et al., 2021). At the leaf level, our study goes much further than what was currently known with a finer hierarchical classification and identification of new signatures SNPs.

A first notice that jumps out is that the tree is that internal nodes are numerous, relatively wide as compared to its depth. This relates with the fact that the means and standard deviation of the differences by sublineage are often large and similar between them, and that the part at > 95% is often null or low, while the number of shared variants is frequently high. In our opinion, these various remarks suggest that diversification *i.e*. transmission was pervasive in the beginning of L5 history but that it diminished in later evolutionary periods. In addition, this ancient diversity could still be underestimated due to under-sequencing of the geographic regions concerned by this lineage. In other words, we would be dealing with a long history for this lineage, which, because of a current disappearance context, would be hidden and would pass “*under the radar*”.

Regarding the genetic events having occurred along this diversification, we can note that there are quite few IS*6110* insertions, RD, and missing genes. A notable exception to the little role of IS*6110* is its insertion between spacers 30 and 35 in CRISPR locus, which is a feature of the L5.1. This overall lack of L5 genomic dynamism could contribute to its disappearance, likely because the bacillus is too slow to adapt and because the long-lasting cross-talk between its host served the host resistance. Hence, the phylogeny described in this paper is bound to evolve, especially since L5 is severely undersampled for the time-being as compared to other lineages.

We now want to remind of the provenance of samples and potential associated biases. L5-caused tuberculosis represents an important part of all tuberculosis in some important regions from Africa such as Nigeria, Benin, Cameroun, Ghana *i.e*. in countries bordering the gulf of Guinea (Coscolla et al., 2021; Gehre et al., 2013; Gygli et al., 2019; Osei-Wusu et al., 2021; Sanoussi et al., 2021; Silva et al., 2022). But in Togo for example, the data of L5-L6 prevalence remains completely unknown. In their 2013 paper describing the phylogeography of L5 in Benin, Nigeria, and Sierra Leone and based on spoligotyping signatures, an attempt to describe 10 L5 sublineages, L5.1 to 5.10 was done, however without whole-genome sequencing approach (Gehre et al., 2013). Actually, the landscape of CRISPR evolution in L5-L6 is complex; from one hand, some sublineages can indeed be described, such as the new L5.3 (new), that bears a yet unknown and rare specific 43 spacers spoligotype (674077717777071). The former L5.3 (Coscolla et al., 2021) is now L5.2.3 and should not considered as a subbranch of L5.2 (new). From another hand, the 43 spacer signature is not always sufficient to assign definitively all other sublineages, although many sublineages can be defined through specific spoligotyping signatures (see **Supplementary Material 3**). We provide in this article a list of 45 SNPs that allows to resolve the phylogenetic tree of L5 with 3 main branches (L5.1, L5.2, L5.3 (new)). In some cases we link specific IS*6110* insertion/deletion events with some particular sublineages emergence such as the loss of spacers 21-24 in relation to the loss of IS*6110* in L5.1. Our result improves the former classification obtained by Ates et *al*. and by Coscolla *et al*. (Ates et al., 2018; Coscolla et al., 2021). Still, a few strains remain difficult to classify (**Figure 10**).

Our study still suffers from some limitations: when genomes are present in databases, the geographical origin of the samples is often missing so that these data are of poor interest for phylogeography. Moreover, data are either absent or scarce for many countries of Africa, that still suffer from undescribed tuberculosis linked to L5 or L6. A striking example is given by Nigeria, a 200 million people country that has only two MTBC publicly available WGS in the NCBI genome database (search done in July 2022). We also identified a trend for specific L5.1 and L5.2 phylogeographic distributions that remain to be confirmed by extensive recruitment in central Africa. Whether this is caused by historical, environmental factors or human genetics factors is however difficult to say. It is commonly accepted that MTBC and its host developed under strong sympatry. This is however based on few facts as research studies trying to link human genetics predisposition to bacterial genetics remain rare (Gagneux et al., 2006).

Long-read sequencing on three geographically distant isolates of L5 suggested that using H37Rv as a reference genome is limiting and showed that the pangenome of L5 could be larger than expected previously (Sanoussi et al., 2021). Preliminary results using *de novo* alignment using Roary indeed suggests that some L5 may harbor specific genes that are not found in H37Rv (results not shown) (Page et al., 2015). However the purpose of this study was not to focus on the pangenome, a challenge that will require more L5 to be sequenced, but rather to focus on SNPs analysis, CRISPR locus structure, classification and phylogeography of L5. The most specific genes found to be absent in L5 and present in H37Rv are Rv1979c and Rv2478c (results not shown).

Recently, an important paper by Kerner *et al*. studied ancient DNA (aDNA) of Europeans to look for the time-distribution of an homozygous P1104A allele on the tyrosine kinase TYK2 gene, an allele that has been found to be associated to an increased susceptibility to tuberculosis in humans (Kerner et al., 2021). Using a collection of more than 1000 ancient human DNA, they suggest that there was an emergence of this allele around 30,000 years ago in west Eurasia, followed by a negative selection that started around the bronze age in Europe (Kerner et al., 2021). These data may also apply to Africa, but DNA from african human populations, *a fortiori* aDNA are less studied for the time-being (Choudhury et al., 2020). A recent study performed in Ghana suggested that there could be a strong anthropological link between the patient group (in this case the Ewe) and the likelihood to have a tuberculosis caused by *M. africanum* (Asante-Poku et al., 2015). Another recent study suggests that patients infected with L5-L6 versus L4 possess distinct intestinal microbiota (Namasivayam et al., 2020).

An interesting but speculative issue concerns the historical timeframe and geographical area of emergence of L5 in a wider evolutionary context. Recently, a new phylogroup, named as MTBAP for “MTB-associated phylotype” was defined, as the “missing link” between a likely environmental ancestor and a specific host-adapted, professional pathogen mycobacteria that includes recently described new species such as *M. decipiens, M. shinjukuense, M. lacus and M. riyadhense* (Sapriel and Brosch, 2019). It is known for infectious diseases agents that, finding the center of diversity, or center of emergence, is not always simple, due to the importance of passed migratory events that may have blurred the original picture (Coscolla et al., 2021; Menardo et al., 2019). However slow-growing pathogens like the MTBC bacilli, living an intra-cellular life-style, that spread through aerosols and airways, that are pathogenic, are more likely to have left long-lasting scars and traces in animals and/or in human populations than rapid, extra-cellular, highly pathogenic pathogens, that are transmitted by an oro-fecal way, and that are more rapidly swept by new variants (Achtman, 2016). In this sense, L5 (and also L6) show an highly geographically-constrained patterns of interest: West of West-africa for L6 and Guinea gulf and central Africa for L5. Considering also the presence of L1 in East-Africa, likely to find its origin in South-Asia (Menardo et al., 2021), and the assumption that tuberculosis was a human-driven rather than a cattle-driven disease (Brosch et al., 2002), one must however consider with great attention the recent results obtained by Campbell and Ranciaro on lactase persistence in human beings in relation to the demographic history of pastoralists (Campbell and Ranciaro, 2021) (**Figure 11**). Whether lactase persistence in humans could be linked to L1-L4-L5-L6 MTBC lineages emergence in relation to cattle domestication remains an open question. Lactase persistence is linked both to animal herding and milk consumption. It has left traces of selection in humans and prevalence of the various adaptation alleles are also linked to the demographic histories of pastoralists population, in relation to climate change (Campbell and Ranciaro, 2021) unless gut microbial composition could compensate as in some pastorialist populations (Campbell and Ranciaro, 2021). This gut microbiota composition could also be associated to differential susceptibility to different lung infection agents (Namasivayam et al., 2020). Whether this is a cause or a consequence of infection remains to be studied, however an indirect link may exist between a nomadic pastoralist and a certain type of lung infections on one hand, and in another hand, a sedentary agricultural lifestyle with other lung infection types. Related to this topic, a surprisingly congruent geographical pattern is found between the presumed time-frame of emergence and area of migration of the two main *Bos* species, *Bos taurus* and *Bos indicus*, and the geographical distribution of L1, L4, L5, L6 in Africa (Campbell and Ranciaro, 2021).

**Figure 11:**
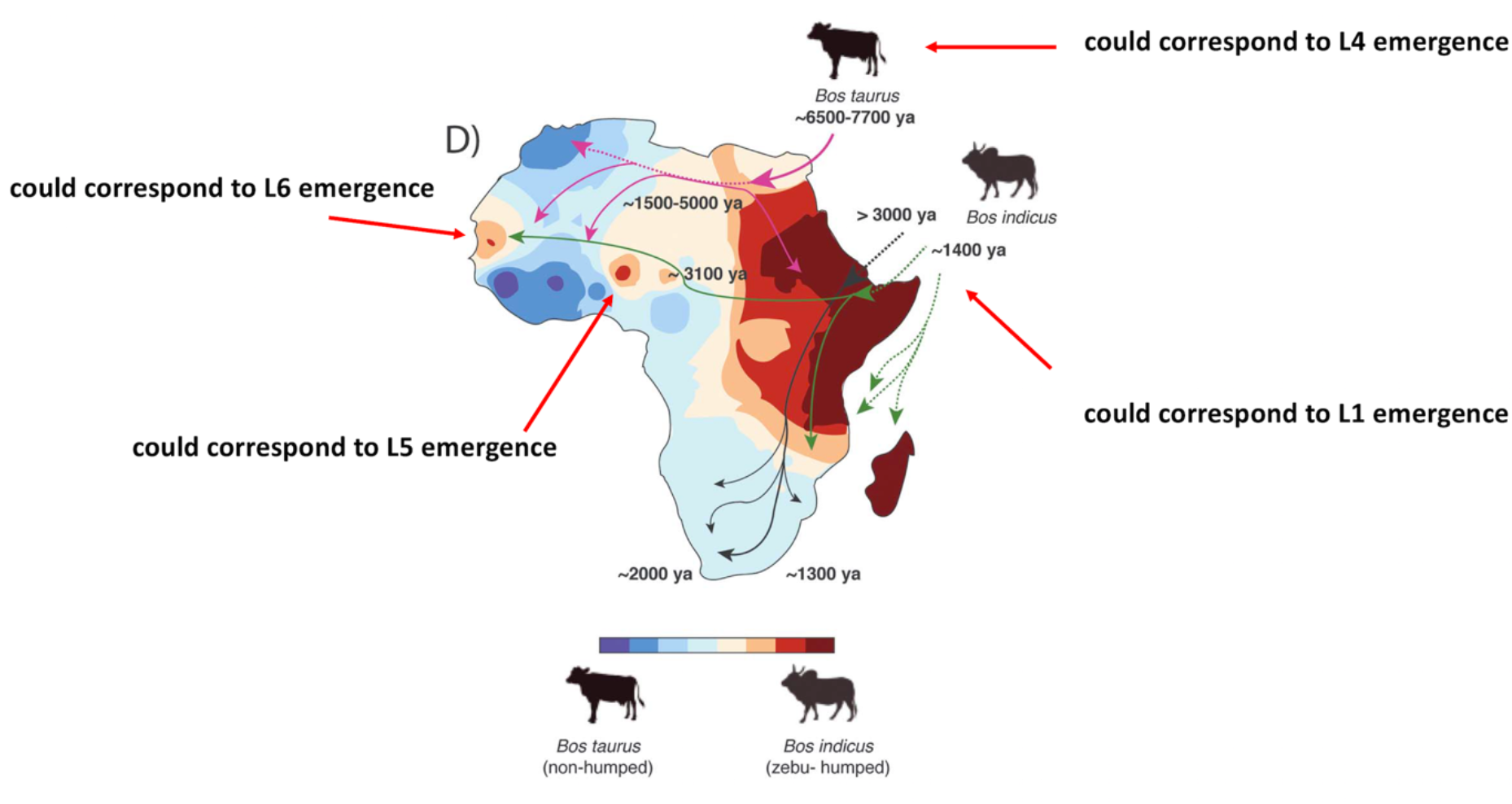
Hypothesis of emergence of L1-L5-L6 (adapted from Campbell and Ranciaro 2021, reproduced with courtesy of the author)

We suggest that an MTBC lineage (L4) could have emerged in relation to a North-african *Bos taurus* domestication, and hence, would be found only first in North of Africa and by extension elsewhere in Africa, contemporary to the recent neolithic expansion (Bantu agricultural expansion) (blue color in **Figure11**), an expansion that would have finally led to a specialized human pathogen. Conversely, we suggest that the other L1-L5-L6 MTBC lineages could have been related to *Bos indicus* domestication and introduction into Africa via East-Africa and the arabian peninsula (red color in **Figure 11**). It is quite puzzling, as shown in **Figure 11** that the two isolated red spots, indicator of *Bos indicus* introduction events found in West Africa quite well correspond and can be superimposed to L5 and L6 geographic distribution (respectively Cameroun-Nigeria and Senegal-Gambia) and could coincide with an early (3000 years ago) or late (>1400 years ago) arrival of *Bos indicus* from the Arabic peninsula, first via Eastern Africa, and later arrival in central Africa and even later in West Africa. Such a scenario also fits with the recent discovery that *M. bovis* also emerged in Africa with nomad pastoralists having domesticated different cattle in the near-east or in the Indus valley (Loiseau et al., 2020). Conversely the L4 sublineages could have spread from West Africa to Central and South Africa following the Bantu speakers expansion (Patin et al., 2017).

Hence, tuberculosis could either have initially been sitting in Africa since paleolithic times in humns, or alternatively have been introduced into Africa many times through various animal or human migrations routes in relation to the Neolithic transition, to human slave trade, and possibly via South Asia and the Arabic peninsula (Comas et al., 2013; Menardo et al., 2021), a complex history that remains to be better understood. L6 emergence is posterior to L5 emergence and the fact that L5 and L6 are respectively more prone to be found in human (L5) or in animal (L6), possibly argues of a potentially long duration of interactions of their last common ancestor with animals and potential co-infection and complex cross-talk history of tuberculosis in Africa (Perrin, 2015). Moreover in Africa, as already mentioned, the “*cattle before crops*” model for food production is indeed very different from anywhere else in the world (Manning, 2010). The neolithization process should also be understood differently in Africa and in Europ, because of a very different history and a deep anthropological significance (Manning, 2010).

To conclude, we believe that it is very likely that the early history of tuberculosis caused by *M. africanum*, is intimately linked to local processes that are linking demographical, anthropological, and ecological factors, including climate-change, that shaped, throughout the millenias, today’s Africa MTBC bacilli population structure, with intricate relationships between human and animal life-styles, from hunter-gatherer, pastoral, agro-pastoral, and more recently urban life-styles. In Africa, an accurate reconstruction of the global remote and/or recent tuberculosis history remains intimately linked to an expansion of numerous new local epidemiological and evolutionary WGS studies. Nevertheless, in this article, we have provided a more precise picture of some of the evolutionary steps of L5 diversification using for the largest L5 sample and a specific tool easing the exploration of all genetic modifications along MTBC history, *TB-Annotator* (v.1, 15900 genomes and v.2 90000 genomes). However, most of the functional consequences of the identified genetic events characterized so far, still need to be studied in more detail to grasp the virulence evolution of all these lineages.

## List of Figures and tables

**Figure 1**: General algorithm of the TB-Annotator

**Figure 2**: General overview of the L5 phylogeny

**Figure 3**: Existing phylogeny (Coscolla et al., 2021) reference naming and transient naming of new clades

**Figure 4**: New phylogeny with hierarchical level naming

**Figure 5**: 5.3 sublineage

**Figure 6**: 5.2.1 and 5.2.4 sublineages

**Figure 7**: L5.2.2 sublineage or L5.2 in (Coscolla et al., 2021)

**Figure 8**: 5.2.3 sublineage

**Figure 9**: L5.1 and L5.2 geographical distribution as found in this study

**Figure 10**: A final phylogeny of Lineage 5 with SNP barcode proposal

**Figure 11**: Hypothesis of emergence of L1-L5-L6 and L4 (adapted from Campbell et al. 2021, reproduced with courtesy of the author)

**Table 1**: conversion table between the reference nomenclature and our new naming

**Supplementary Material 1** Table 2: List of 45 SNPs, Lineage information at internal of leaf node, detail information on SNPs assessed and complementary informations (xls file)

**Supplementary Material 2** A thorough classification of the L5 Sub-lineages (pdf. File, 10 pages).

**Supplementary Material 3**: Table 3: Accession List of genomes assessed, CRISPR-builder reconstructed spoligotype, countries of origin of the samples, miscellaneous information on CRISPR, comparison of classification results obtained between TB-Profiler lineage assignment, automatized spoligotype recorded in TB-Profiler, Final list of all spoligotypes (L5-spoligo-all), results of Ates et al. reanalyzed (xls. File, 47 pages)

**Supplementary Material 4**: Table 4: Accession List of genomes assessed, with all different L5 sublineages SNPs specificity search in TB-Annotator v1 (15900 genomes) (xls. File, 45 pages)

**Supplementary Material 5**: Two phylogenetical trees built using RaXML with or without PE-PPE gene variants on a subset of samples. (pdf. File, 3 pages).

## Author statements

### 4.1 Author contributions

Conceptualization: CG, GS, GR, CS; data curation: CG, GS, CS, MRS, BMM; Formal analysis : CG,CS, MRS; Funding acquisition: CS, EC, CG; Investigation: CS, GR, CG, MRS, KL; Methodology: GS,GR,CG, MRS; Project Administration: TP,MRS,JD; Resources: EC;JD,CG,TP; Software: GS,KL, CG; Supervision:CS,CG, EC; Validation: CS,KL,MRS; Visualization: CS,CG; Writing original. Manuscript; CG,CS,GR, MRS; Writing-review and editing: CG, CS, MRS, GR.

### 4.2 Conflicts of interest

The author(s) declare that there are no conflicts of interest

### 4.3 Funding information

This study was supported by PTDF (Petroleum Technology Development Trust) of Nigeria through a PhD grant attributed to MRS, by KNCV Nigeria (TP), by Campus-France through benchfees for MRS, by the UMR1137 IAME INSERM-Université Paris-City research Unit, and by the APHP (Assistance Publique des Hôpitaux de Paris) through the associated French NRL for TB. This project is part of MRS PhD (phylodynamics of MTBC with a specific focus on L5-L6 lineages).

### 4.4 Ethical approval

Public health action taken as a result of notification and surveillance is one of the tasks of the *Service de mycobactériologie spécialisée et de référence* of the Bichat Hospital in Paris, as a French TB NRL -associated laboratory, with identical missions as the one of NRL which provide a mandate and legislative basis to undertake necessary follow-up. Part of this follow-up is identification of epidemiological and molecular links between cases and research. This study is carried out under this framework, and as such explicit ethical approval was not mandatory at start working on archive material. However a prospective study, whose results are going to be reported elsewhere is currently under approval at the Nigerian Federal NRL ethical committee. Patient data are not traceable by any other third part than by the NRL. This research is done in the framework of the PhD thesis of MRS, grantee from the PTDF (Petroleum Technology Development Fund) of Nigeria.

### 4.5 Consent for publication

All authors read and approved the final version of this article.

### 4.6 Acknowledgements

This study is dedicated to Dr. Lovett Lawson, Nigeria STOP-TB officer, who passed away due to Covid19 in 2020. We are grateful to Barbara Molina-Moya, Jose Dominguez and Luis Cuevas for providing us with DNA archives of L5-L6 from the previous EDCTP Collaborative study led by Pr. Luis Cuevas and Pr. Lovett Lawson. We are grateful to Dr. Jody Phelan, from the LSTM for providing us with an updated Spoligotyping list with associated SRA accession number. MRS is a PhD fellow from Nigeria supported by the Petroleum Technology Development Fund from Nigeria (PTDF) which is warmly acknowledged; Campus France is also acknowledged for his support and the IAME research Unit, UMR1137, and its director, Pr. Erik Denamur, for welcoming and supporting MRS.

## Footnotes

1 In the same vein, the more one advances in the lineage, the more the spoligotype is in confetis.

## Notes

### Competing Interest Statement

The authors have declared no competing interest.

### Summary of Updates

suppression of TB-Annotator URL address in the new version

