## Supplementary material 2 for "An updated evolutionary history and taxonomy of *Mycobacterium tuberculosis* lineage 5, also called *M. africanum*" (from another version of this preprint: version 1, 2022-11-21)

### A thorough classification of the L5 sub-lineages

#### 1 Investigating the 5.1 lineage

##### 1.1 The 5.1.1. case

This sublineage is constituted by two sublineages: the new L5111 and the L5112 that contains 5.1.1., 5.1.2., and 5.1.3.

**The 5111 clade** L5.1.1.1 is constituted by 15 isolates, cf. **Supplementary Fig.1**. It is defined by C2135816G and 7 other exclusive SNP variants (0 at > 95%). There is an average distance of 432 SNPs between these isolates (with a standard deviation of 186).

The structure of 5.1.1.1 lineage depicted in **Supplementary Fig.1** is obvious: two clades, the second one containing two sub-lineages.

The first clade 51111 is constituted by two isolates, namely: ERR3170415 and ERR3170414 from Ghana. These two isolates are quite different, having a distance of 121 SNPs. They are far from the other isolates, presenting 205 exclusive variants (0 > 95%), one of them being C42625A. Surprisingly, they have 5 exclusive IS*6110*s, at the following positions: 1541099, 1934645, 2199408, 2634166, and 4198463. They additionally share 4 exclusive regions of deletion: 844720-845055, 1188589-1188650, 3079164-3079309, and 4211161-4212325.

The second clade 51112 has 12 isolates; their mean difference is of 347 SNPs, with a std of 166. It has 3 exclusive variants (0 > 95%), one of them being C821895T. It is composed by two subclades.

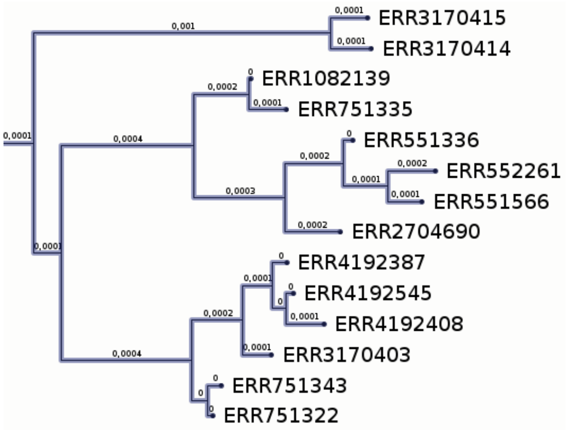


**Supplementary Fig.1**: 5.1.1.1 sublineage

The first one 511121 at the top of the clade is constituted by 6 isolates that have 21 exclusive variants (0 > 95%), e.g., G36400C. They are at a mean distance of 242 SNPs (87 std).

The second one, 511122, contains 6 isolates too and is defined by C81787A. They share 102 exclusive variants (0 > 95%), and they are at a mean distance of 123 (51 std).

**The 5112 clade** 5.1.1.2 is the last common ancestor of 5112232 (5.1.1), 511222 (5.1.2), 51121 (5.1.3), and the new 511221, 5112231 and 511222 lineages.

It is constituted by 193 isolates that share 18 exclusive SNP variants (16 more at 95%), one of them being G141375A. They all have a deletion of Rv1522c. The mean distance is of 435 SNP, with a standard deviation of 98.

L51121 (5.1.3, Figure 10) is constituted by 18 isolates that are defined by G191218C. They share 19 variants in exclusivity (0 > 95%), and have a mean difference of 381 SNPs (std of 77). And as can be seen in **Supplementary Fig.2**, the 51121 lineage can be subdivided in two sublineages.

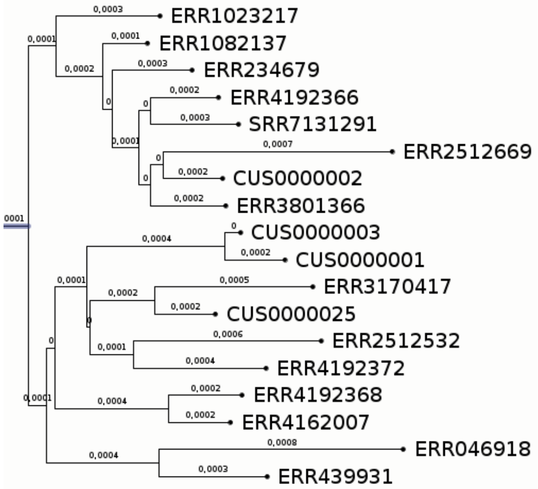


**Supplementary Fig.2**: 5.1.1.2.1 sublineage

At the top, 8 isolates are defined by 15 exclusive variants (0 > 95%), one of them being C70090A. They have an average of 270 (73), and are denoted by 511211. The remainder L511212 of the tree, however, does not present exclusive variants, and it may not be phylogenetically consistent.

L51122 is the last common ancestor of 5112232 (511 in Coscolla et al. [7]), 511224 (512 in Coscolla et al. [7]), plus various small new lineages. It is constituted by 175 isolates, that all share C2081285T. They have 3 exclusive variants, and 12 variants larger than 95% of exclusivity. The mean difference is of 423 SNPs with a standard deviation of 100. Its sublineages are described below.

L511221 is defined by G106903C. It contains 15 isolates, which have 5 exclusive variants (0 > 95%). The average difference is of 373 SNPs with a standard deviation of 76.

L5112211 is constituted by 6 strains, as depicted in **Supplementary Fig.3**. It is defined by C7508T, and it has 30 exclusive variants (0 at 95%). The mean distance is of 242 SNPs with a standard deviation of 73.

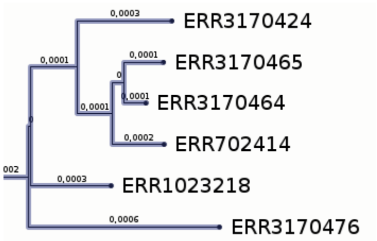


**Supplementary Fig.3**: 5.1.1.2.2.1.1 sublineage

L5112212 is constituted by 9 strains, as shown in Figure 12. These isolates share 16 exclusive variants (0 at 95%), such as T164818C. They have a mean distance of 376 SNPs with a standard deviation of 60.

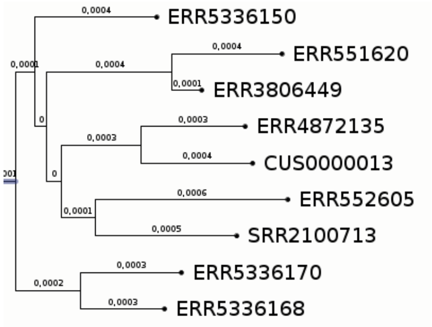


**Supplementary Fig.4**: 5.1.1.2.2.1.2 sublineage

L511222 (512 in Coscolla et al. [7]) as shown in **Supplementary Fig.4**is constituted by 29 isolates which share 29 exclusive variants (6 > 95%), like C71636T. Its mean difference is of 333 for a standard deviation of 85. The 11 isolates at the top of the L511222 tree define a sublineage denoted by L5112221. They are constituted by 12 exclusive variants (0 > 95%), one of them being C247157T. They have a mean distance of 317 SNP, with a standard deviation of 114.

The L5112222 at the bottom, with 18 isolates, has 2 exclusive variants (0 > 95%). They are defined by A1230368G, and have a mean of 306 SNP differences (with a standard deviation of 74). L5112222 is constituted by 3 sublineages, whose relationship is not obvious.

The first one is constituted by SRR2101063 (NoAfr), ERR2513725 (unk), ERR1215478 (GHA), ERR439937 (BEN), SRR2100183 (NoAfr), ERR3170471(GHA), ERR1203060 (GHA), and ERR1203058 (GHA) (8 isolates, mean: 267, standard deviation 59). It is denoted by L51122221, and only one exclusive variant is shared, namely: A320524C (0 > 95%). The second one, L51122222, is constituted by ERR3170466 (GHA), ERR2514205 (unk), SRR7070014 (unk), ERR1082122 (GHA), and ERR1082123 (GHA). They have 21 exclusive variants, one of them being G1312478A (0 > 95%). Its symmetric difference matrix has an average of 215 SNPs for a standard deviation of 106. The third and last one, L51122223, is constituted by: ERR234199 (GHA), ERR1082119 (GHA), ERR1023224 (GMB), ERR5330907 (unk), and ERR5336146 (unk). It is defined by C280350T, and shares 13 exclusive variants, with a mean difference of 241 for a standard deviation of 112.

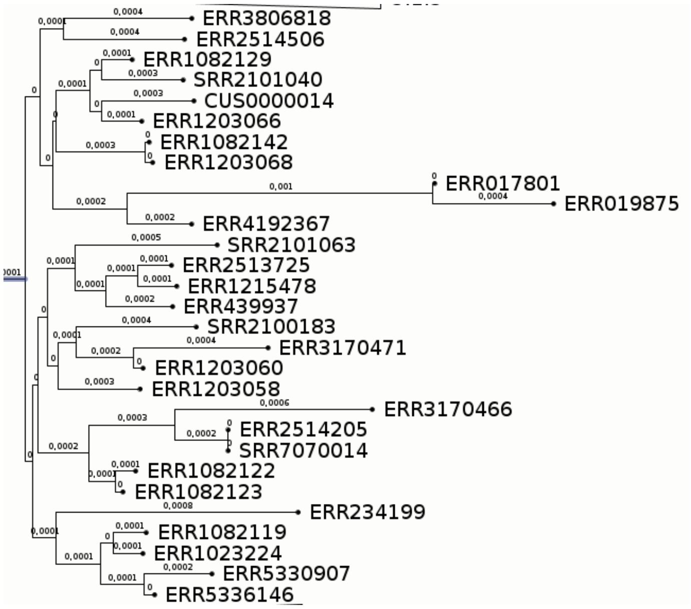


**Supplementary Fig.4**: 5.1.1.2.2.2 sublineage

L511223 is at the origin of L5112231 and L5112232 (L511 according to Coscolla *et al.* [7]). It is defined by C65432A and constituted by 130 strains. It has 14 exclusive variants (37 at > 95%), and a mean distance of 388 SNPs (102 standard deviation).

L5112231 is currently constituted by 3 strains that share 38 exclusive variants (0 variants > 95%), as shown in **Supplementary Fig.5**. They are defined by C145470T, and a total of 38 exclusive variants (0 > 95%). They have a mean distance of 312 SNPs, for a standard deviation of 96.

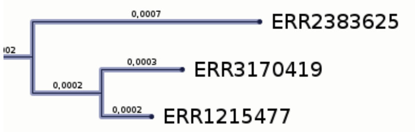


**Supplementary Fig.5**: 5.1.1.2.2.3.1. sublineage

Finally, L5112232 is the L5.1.1 lineage of Coscolla *et al.* [7]. It is constituted by 127 isolates that have 21 exclusive variants (30 > 95%). They can be defined for instance by C11846G, and they present a mean distance of 385 SNPs (with a standard deviation of 102). This tree is not shown.

##### 1.2 The 5.1.2. case

This lineage is constituted by two sublineages that have already been defined by Coscolla *et al* [7].

L5.1.2.1 (5.1.4 in Coscolla *et al.*) is constituted by 47 isolates (cf. **Supplementary Fig.6**) that are defined by T221767C. They have 20 exclusive SNP variants (6 more at > 95%). They have an average distance of 372 SNP with a standard deviation of 99.

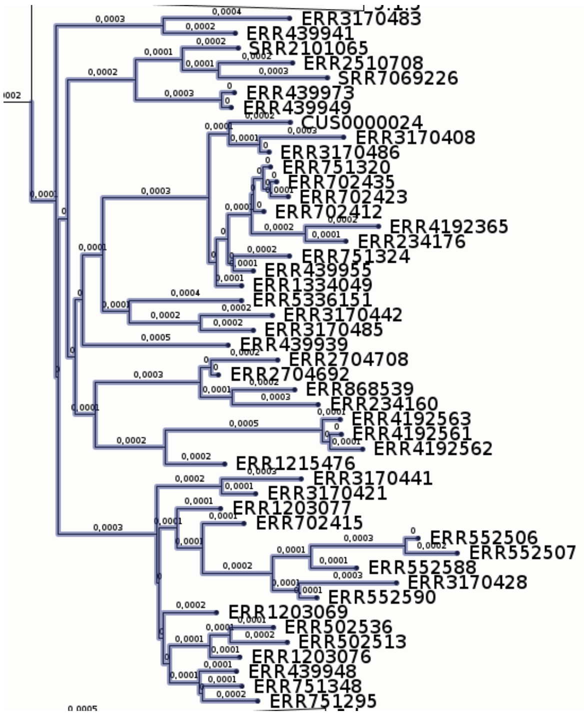


**Supplementary Fig.6**: 5.1.2.1 sublineage

Two genomes appear at a basal position in this lineage, namely: ERR3170483 and ERR439941. They are somewhat particular, sharing 72 exclusive variants (0 > 95%) plus one exclusive region of deletion (3741286-3741419). For instance, A255406G is exclusive. Furthermore they are quite distant, as 267 SNPs separate them. They can be the apparent part of a clade to be completed, which we will note 51211.

The 45 remaining isolates constitute a clade too, having 2 exclusive SNPs (G1365878A and C3855611T), plus 20 variants of exclusivity larger than 95%. The mean distance is equal to 368, for a standard deviation of 101. It will be denoted as 51212. This lineage seems to be constituted by 3 clades, whose phylogenetic resolution is not obvious.

The first clade, namely 512121, is constituted by 5 isolates with a mean difference of 245 (80). They share 40 exclusive variants, (0 > 95%), T3471G among other things. An IS*6110* is quasi exclusive too, being present in these 5 isolates plus another one outside the selection. It is at position 2266177. A region of deletion is specific too to this clade, namely at position 2854958- 2855299.

The second clade, namely 512122, has 24 isolates. It is defined by 4 exclusive variants, among which C1533697T. Two other variants are exclusive at > 95%. They are at average distance of 316 SNPs, with a standard deviation of 117.

The third clade at the bottom of the tree of **Supplementary Fig.6** is 512123, defined by G115073A. It is constituted by 16 isolates having 59 exclusive variants (0 > 95%). Their mean is of 245 SNPs, for a standard deviation of 68.

L5.1.2.2 (5.1.5) is constituted by 18 isolates (see **Supplementary Fig.7**), and is defined by C24663G. They share 78 exclusive variants (0 at > 95%). Two IS*6110* are of particular interest here, namely the one at position 3486353, which is exclusive to this lineage and present in 17 isolates (ERR439940 does not have this insertion), and the one at position 482905, present everywhere here and in 4 other isolates. The region between positions 968974 and 969052 is removed in all the isolates, except for ERR552427. Finally, the mean distance is of 260 SNP, with a standard deviation of 98.

5.1.2.2 contains one sublineage at the top of the tree of **Supplementary Fig.7**, called 51221, and which is defined by C60707T. 95 variants are exclusive (0 > 95%) to these 7 strains. They shared too a small missing region of difference, namely at position 2970579-2970661. Their distance is low, with a mean of 74 SNPs (std 25), but we still do not face here an epidemic event. The 11 isolates left are for the time-being not further classified in a more phylogenetically significant clade 51222.

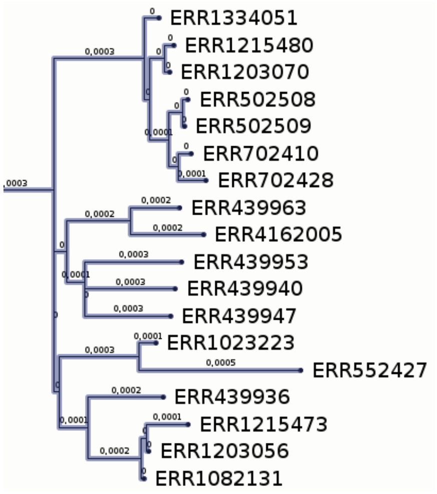


**Supplementary Fig.7**: 5.1.2.2 lineage

#### 2 Investigating the 5.2 lineage

##### The 5.2.2. case

This clade is constituted by two subfamilies, (cf. **Figure 7 in main manuscript**). At its top, we find 5221, which is constituted by 17 isolates. It has two exclusive variants (0 > 95%), such as G353427T. Their mean distance is of 326, for a standard deviation of 207. This clade may be not perfectly defined. The lineage 5222 at the bottom is constituted by 19 isolates. They share 51 exclusive variants (0 > 95%), one of them being T5124C. They have an exclusive IS*6110* at position 3760397. Finally, they are at an average distance of 325 SNPs with a standard deviation equal to 102.

##### The 5.2.3. case

There is a first clade at the top of the tree of Lineage 5.2.3, (cf. **Figure 8 in main manuscript**). This 5231 sublineage is constituted by 14 isolates that share 44 exclusive variants (0 > 95%). They are defined by C29706T. They share a large missing region of difference, from position 1683109 to 1717671 (from Rv1492 to Rv1522), and they have a mean difference of 276 SNPs (std: 86).

The clade at the bottom is constituted by 15 isolates, it is denoted by 5232. It is defined by G904244C and has 13 exclusive variants (0 > 95%). The mean distance is of 431, with standard deviation of 106.

This clade is subdivided into 2 subfamilies. The first one at the top, 52321, is constituted by 6 isolates that are defined by G258120A. They share 10 exclusive variants (0 > 95%), and have a mean of 416 SNPs (std: 123). At the bottom, 52322 has 9 isolates at an average distance of 345 SNPs (100). They are defined by A192438G and present 26 exclusive variants (0 > 95%
