## Supplementary material 5 for "An updated evolutionary history and taxonomy of *Mycobacterium tuberculosis* lineage 5, also called *M. africanum*" (from another version of this preprint: version 1, 2022-11-21)

#### RAxML_bestTree.withPE-PPE.pdf built on a subset
#### Supplementary File 5.1

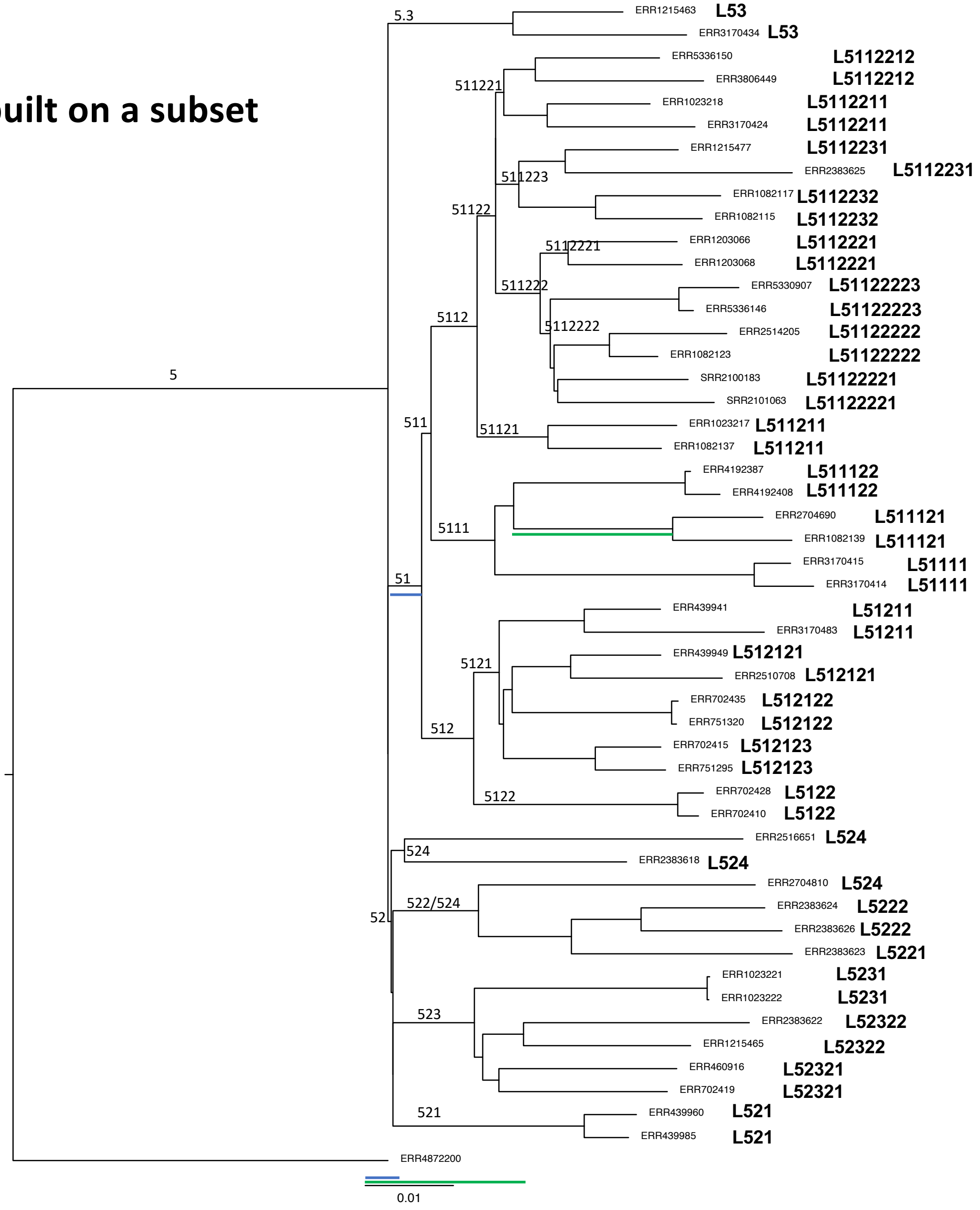

### RAxML_bestTree.withoutPE-PPE.pdf built on a subset
### Supplementary File 5.2

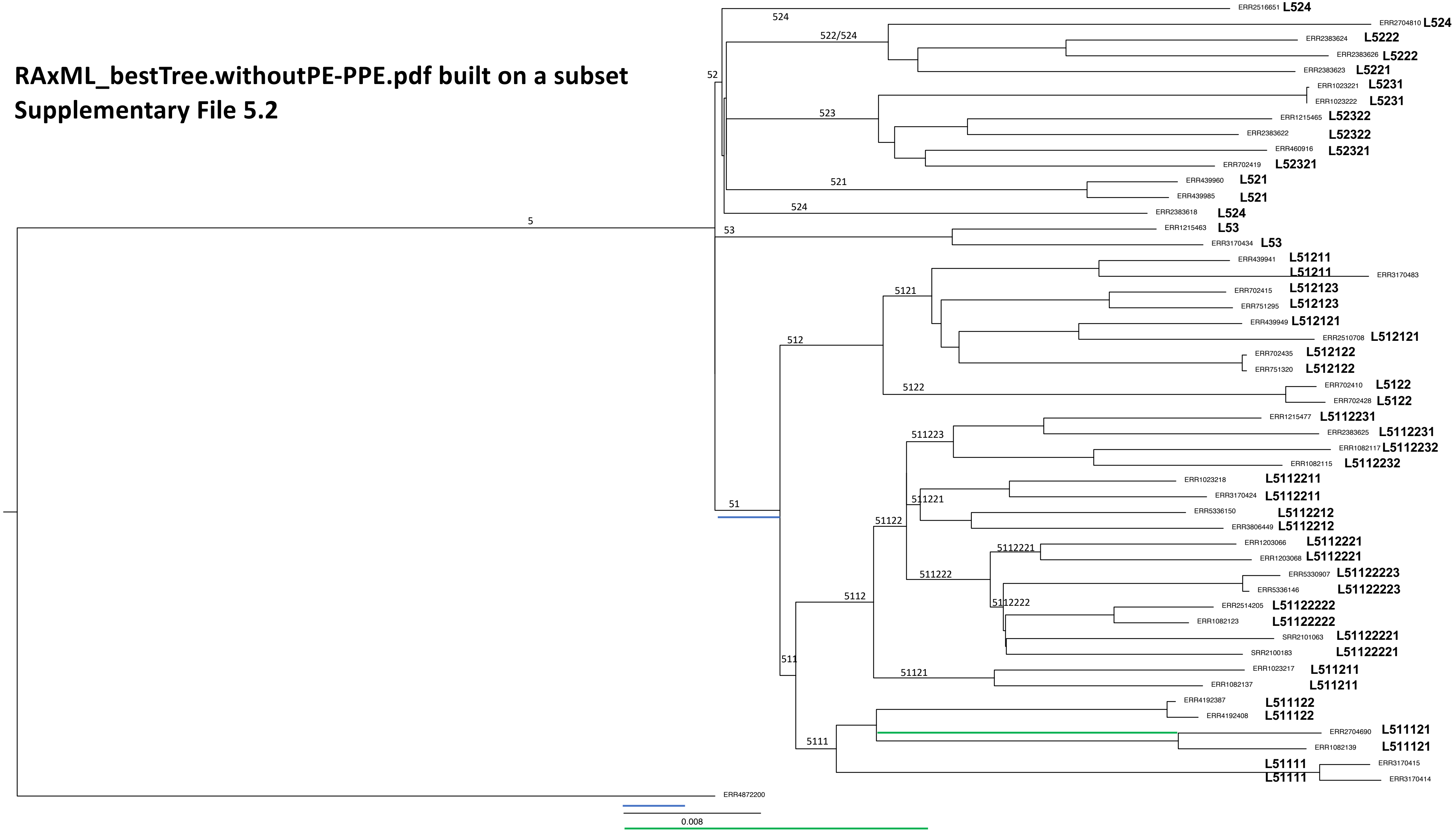

**RAxML_built Tree with PE-PPE and RAxML_built Tree without PE-PPE, same but trees displayed on ITOL, see ref. below Supplementary File 5.3**

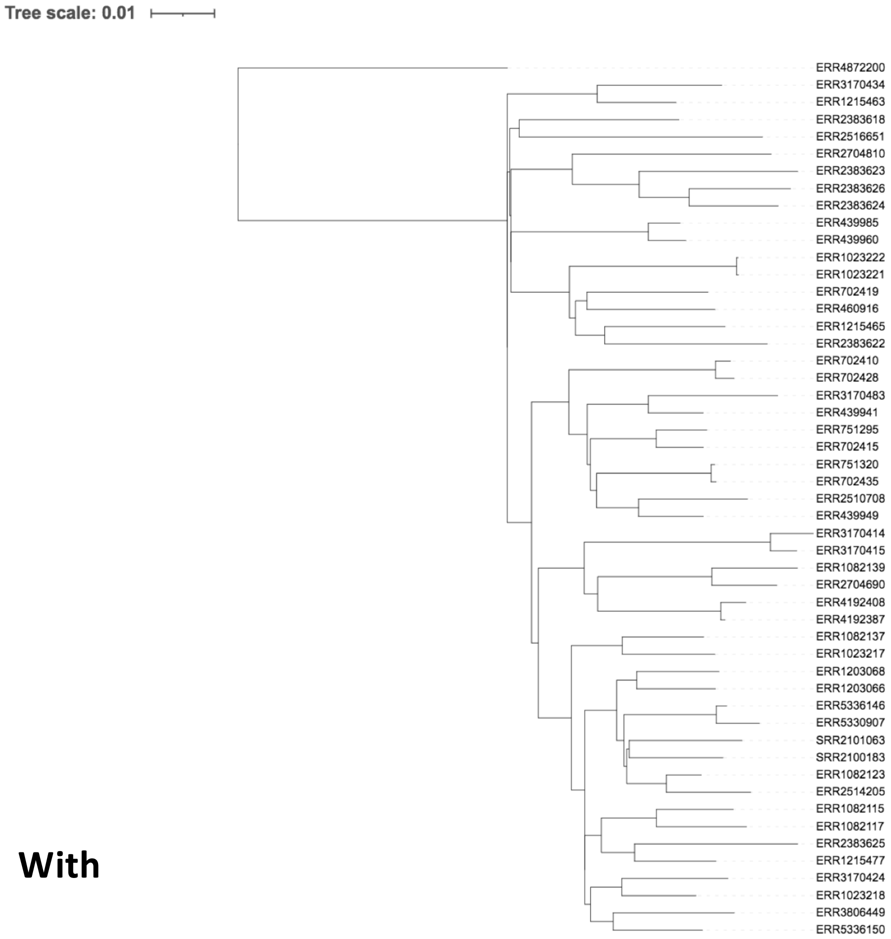


**With**

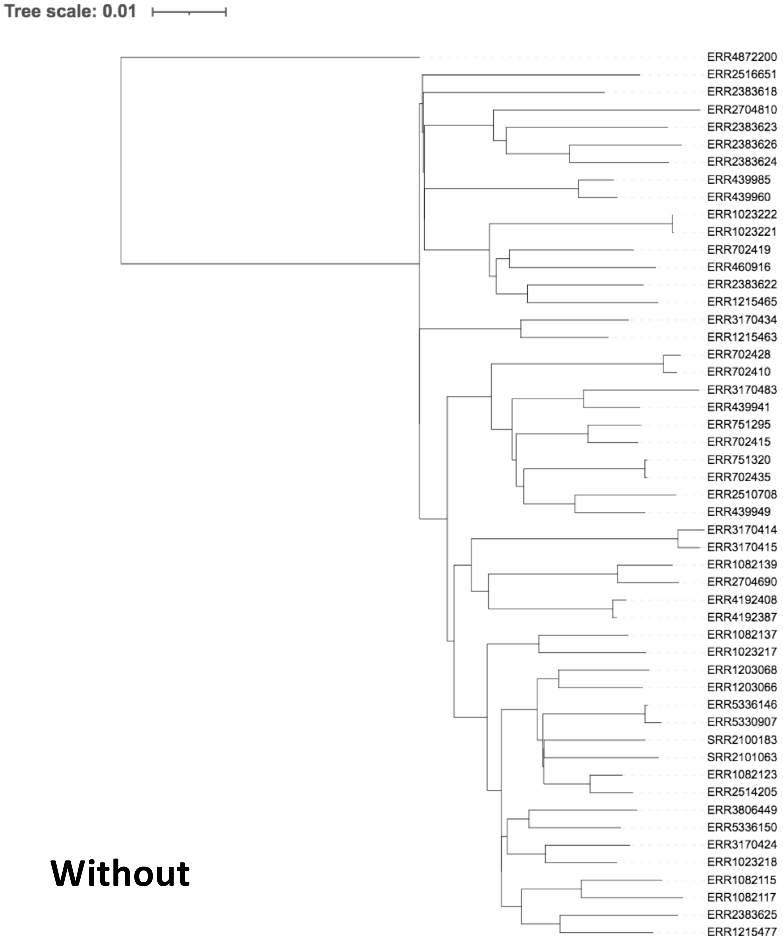


**Without**

Letunic and Bork (2021) *Nucleic Acids Res* [doi: 10.1093/nar/gkab301](https://doi.org/10.1093/nar/gkab301) |
